# Estimating Gene Prevalence with Incomplete Genomes: Application to the Prochlorococcus Pangenome

**DOI:** 10.64898/2026.10.19.760200

**Authors:** Stephen Blaskowski, Louis Goldwater Christie, Yubin Raut, Jed A. Fuhrman, Jacob Bien, E. Virginia Armbrust

## Abstract

Gene prevalence is an essential statistic in population genomics, defined as the proportion of a population carrying at least one ortholog of a given gene family. Common approaches to estimating gene prevalence either assume complete genomes or exclude genomes below a completeness threshold, resulting in biased and high variance estimates that fail to take advantage of all available data. Here we developed a maximum-likelihood framework that explicitly models genome completeness in order to estimate gene prevalence and associated confidence intervals from collections of complete and incomplete genomes. We applied the methodology to marine Prochlorococcus, a globally abundant photosynthetic bacterium for which the majority of available genomes are incomplete. Our methodology produced gene prevalence estimates with associated confidence intervals for a total of 52,500 gene families, more than 4 times the number quantifiable using complete genomes alone. We find a core of 714 genes are likely universal to the Prochlorococcus genus and a substantial majority of 49,828 gene families occur at low prevalence. Our results reveal a Prochlorococcus pangenome dominated by rare flexible genes and provide a general framework for incorporating incomplete genomes into quantitative analyses of microbial genomic diversity.

## Introduction

Individual genomes from the same microbial taxon often diverge significantly in gene content. The ‘pangenome’ is the union of all genes present within a given population and thus represents the full genetic variation and physiological potential available to the population [1, 2, 3]. The pangenome consists of ‘core’ genes common to all genomes and ‘flexible’ or ‘accessory’ genes present in a subset of genomes, and the overall size of the pangenome is hypothesized to reflect the flexibility of a given taxon across different environmental conditions [3]. The marine cyanobacterium *Prochlorococcus* is characterized by an ‘open’ pangenome wherein over half of the genes in any given genome are flexible genes [4, 3]. *Prochlorococcus* is the most abundant photosynthetic organism on Earth [5], and its flexible genes encode functions critical to cell physiology, population fitness, and biogeochemical cycles, including nutrient transporters, cell surface structures, and diverse metabolic pathways [6, 7]. For these and other genes of interest, ‘gene prevalence’ can be quantified as the fraction of a population carrying at least one ortholog copy from a given gene family. Also referred to as gene frequency or gene commonality, [8] gene prevalence defines different categories within the pangenome in addition to core and flexible genes: ‘shell’ genes are found at intermediate prevalence and ‘cloud’ genes are rare, exclusive to small subsets of the population [9]. Thus, gene prevalence reflects complex ecological and evolutionary dynamics including gene gain and loss [10], vertical and horizontal gene transfer [11], predation [12], mutualism [7], and niche partitioning [13]. An accurate quantification of gene prevalence is central to understanding the pangenome of any microbial population.

Previous pangenome studies commonly assumed that input genomes are complete, sometimes imposing quality filters to ensure this assumption [1, 14, 15, 16, 17]. When only complete genomes are analyzed, gene prevalence is simply calculated as the mean of gene presence/absence across the sampled genomes (Figure 1A). However, as Li and Yin demonstrated, the number of detected core genes is reduced by more than half when genomes with an incompleteness as low as 1% are included in the analysis. [18]. This potential problem is compounded by sequencing technologies such as single-cell amplified genomes (SAGs) and metagenome-assembled genomes (MAGs) that commonly return incomplete genomes representing as little as 5% of the genome or less. In this setting, it is impossible to distinguish whether a gene absence can be attributed to a true absence in the genome or to a failure to detect a gene that is actually present (Figure 1B). Some methods have attempted to accommodate genome incompleteness either by lowering the prevalence threshold for categorizing a gene as core [19, 20], or by introducing a more permissive category of an ‘extended core’, commonly defined as genes found in at least 90% of genomes [13]. Other methods have addressed the problem of genome incompleteness more explicitly by estimating the probability that a gene is core based on its presence/absence pattern across genomes [21, 22, 23] or by integrating additional information such as gene synteny to identify core genes [24]. The output of these methods is a categorization of genes as core versus flexible, rather than a continuous estimate of gene prevalence. None of these approaches incorporate information from incomplete genomes into a quantitative estimate of the prevalence of a given gene family in a population of interest.

**Figure 1.**
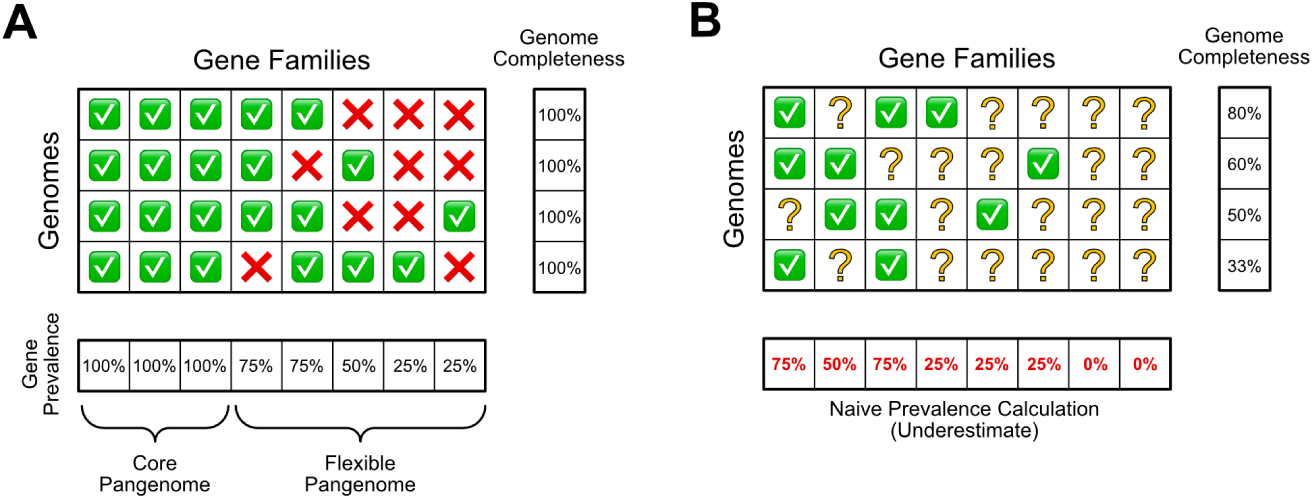
Calculating gene prevalence from gene presence/absence data is nontrivial with incomplete genomes. A) Matrix of gene presence/absence data with complete genomes. In this case gene prevalence can be calculated as a column average. B) Matrix of gene presence/absence data with incomplete genomes. In this case each absence may represent either a true gene absence, or a present gene that was not detected, and thus column averages result in a biased under-estimate of gene prevalence.

In this study we develop a statistical model that explains observed patterns of gene presence and absence as a product of true underlying gene prevalence in combination with the completeness of the genome sequences sampled. We compare different parameter estimation techniques and identify a maximum likelihood estimation approach that faithfully recovers accurate gene prevalence values in simulated datasets. We then apply the methodology to a dataset of *Prochlorococcus*, mostly consisting of incomplete SAGs, and illustrate that integrating incomplete genomes results in higher accuracy estimates of prevalence for a greater number of genes than is possible using complete genomes alone. Finally we compare patterns of gene prevalence within *Prochlorococcus* clade subpopulations, and produce an estimate of global prevalence for every detected gene in the *Prochlorococcus* pangenome.

## Results

We collected a set of 1,099 *Prochlorococcus* genomes, both complete and incomplete, with the goal of estimating the global prevalence of every gene family in the detected pangenome. Here we define ‘gene family’ as a group of orthologous genes, and ‘gene prevalence’ as the proportion of the population carrying at least one ortholog of a gene family. The dataset included 65 closed (complete) isolate genomes, 1,014 single-cell amplified genomes (SAGs), and 20 incomplete draft isolate genomes (Supplementary File 1). Based on CheckM2 score [25], a total of 1,034 of the genomes (94%) were incomplete, and of these over half were less than 60% complete (Supplementary Figure 1), thus precluding use of current methods to estimate gene prevalence despite the size and diversity of the compiled dataset. Incorporating incomplete genomes into prevalence estimations is nontrivial because it is impossible to distinguish truly missing genes from genes not observed due to incomplete sequencing (Figure 1B), and because of this they have historically been excluded from pangenome analysis. However, omitting incomplete genomes discards the majority of available data and eliminates the possibility of estimating prevalence for any gene families that were exclusively detected in incomplete genomes. Therefore, we were motivated to develop a methodology for estimating gene prevalence capable of incorporating incomplete genomes, which are the vast majority of available genomes.

### Model

We developed a statistical model that describes the pattern of gene presence and absence observed in a collection of potentially incomplete genome sequences (Equation 1). The observed presence/absence data are represented by matrix **X** (Figure 1), in which each row *i* represents a genome in the collection, and each column *j* represents a family of orthologous genes. The data is binary, with **X***_ij_* = 1 indicating that at least one ortholog from gene family *j* was detected in genome *i*, and **X***_ij_* = 0 indicating no detection. We modeled each entry **X***_ij_* as the product of two Bernoulli random variables **Y***_ij_* and **Z***_ij_*. Matrix **Y** represents the underlying presence/absence matrix that would result if the true gene content of each genome was known. Matrix **Z** represents the observation matrix, where **Z***_ij_* = 1 indicates that gene *j* was successfully observed in genome *i*. Therefore, every entry **X***_ij_* = 0 is the result of either **Y***_ij_* = 0 or **Z***_ij_* = 0, or both.

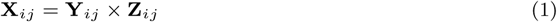

We modeled the presence/absence matrix **Y** as having independent entries **Y***_ij_* ∼ Bernoulli(*π_j_*), such that the probability any particular sampled genome has gene *j* is determined by a population-level prevalence *π_j_*. We modeled the observation matrix **Z** as having independent entries **Z***_ij_* ∼ Bernoulli(*λ_i_*) for some per genome observation probabilities *λ_i_*. These probabilities quantify the completeness of each genome, and in this simplest version of the model, we assume that every gene in each genome has an equal probability of being observed. Given this framing, we next evaluated several different approaches for estimating the *π_j_* prevalence parameters, given the observed presence/absence matrix **X**, and an estimation of the completeness of each genome *λ_i_*.

### Comparison of Parameter Estimation Approaches

We compared the performance of four parameter estimation methods using a simulated dataset of 1,000 genomes and 12,000 gene families. The methodologies included: 1) a naive estimator that calculates gene prevalence estimates as the column means of the observed presence/absence data matrix **X** without adjusting for genome completeness (Figure 1B), 2) a subset estimator that uses the same approach as the naive estimator, but subsets the rows of **X** to only those representing complete genomes (Figure 1A), 3) an unbiased estimator that compensates for missing genes by giving greater weight to gene detection events in less-complete genomes, and 4) a maximum likelihood estimator (MLE) that searches for the prevalence values that maximize the likelihood of the observed presence/absence and genome completeness data (see Methods). The simulated true prevalence values emulate distributions found in real world pangenomes [26, 27, 8], with a majority of genes set as flexible genes with a prevalence that peaks near 0, and a subset of core genes (n=915) with a prevalence equal to 1 (Supplementary Figure 2A). We simulated incompletely sequenced genomes by randomly masking a subset of the true underlying presence values for each genome. The completeness parameters *λ_i_* were drawn from a distribution with 10% chance that *λ_i_* = 1 and 90% chance of being drawn from a uniform distribution (Supplementary Figure 2B).

We evaluated the accuracy of each parameter estimation approach by calculating the mean squared error (MSE) between the simulated true prevalence values of each gene family (*π_j_*) and the estimated prevalence (*π_j_*). As expected, the naive estimation approach underestimated prevalence for almost all gene families (Figure 2A). The subsetting approach avoided the systematic error seen with the naive estimator (Figure 2B), but the estimates exhibited a wider spread than those from either the unbiased estimator (Figure 2C) or the MLE approach (Figure 2D), as well as wider confidence intervals, as a result of using a fraction of the available data. Overall, the MLE approach resulted in the lowest error across all gene prevalence categories, except for core genes, for which the subset estimation approach resulted in perfect accuracy (Figure 2E).

**Figure 2.**
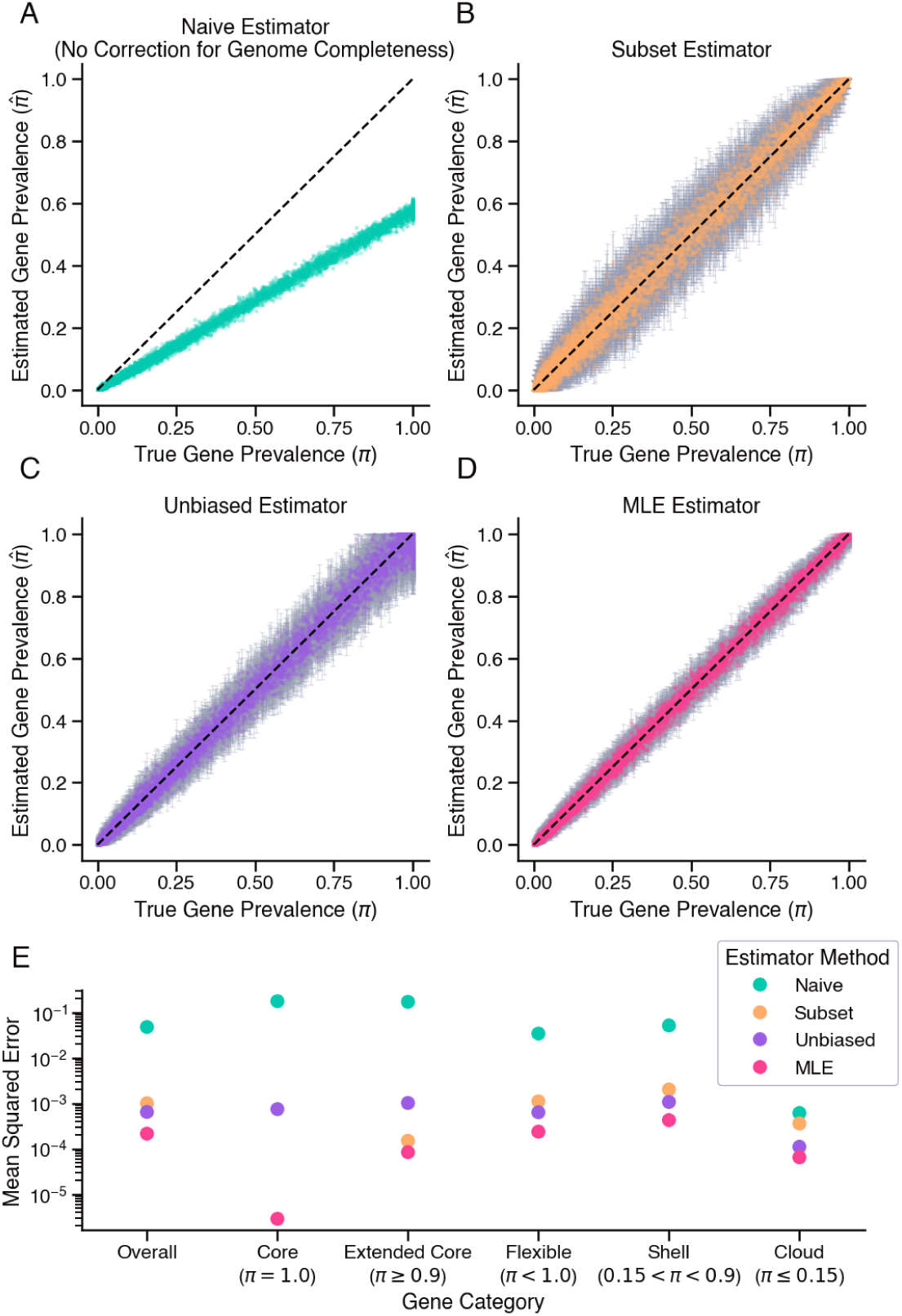
Maximum Likelihood Estimate (MLE) method exhibits lowest error in recovering true gene prevalence values in simulated data. A-D) Model-predicted versus true gene prevalence as predicted by A) naive method that does not account for genome completeness, B) methodology that considers only the subset of complete genomes, with 95% confidence intervals, C) unbiased estimator based on Bernoulli model with 95% confidence intervals, and D) MLE estimator with 95% confidence intervals. Black dotted line indicates perfect predictions. E) Mean squared error exhibited by each method, subset by different gene categories.

We next developed and compared methodologies for the construction of confidence intervals associated with the prevalence parameters *π_j_*(see Methods). For the subset and unbiased estimators, the confidence intervals were based on the estimator’s variance and were constructed via a Gaussian approximation. For the MLE we used likelihood ratio confidence intervals. We evaluated the performance of these inferential methodologies across different confidence levels and gene prevalence categories (Supplementary Figure 3). At every confidence level, the Gaussian approximation approach used in combination with the unbiased estimator resulted in wider confidence intervals than the likelihood ratio methodology used in combination with the MLE approach (Supplementary Figure 3A). At the 95% confidence level, this trend held true across all gene prevalence categories, with a mean confidence interval of about 0.1 for the unbiased estimates of extended core genes (Supplementary Figure 3B). This means that for this subset of genes the confidence intervals were on average as wide as the prevalence interval of the category itself. In contrast, the confidence intervals for the MLE approach never exceeded 0.1 in any category and averaged around 0.03 for extended core genes. We also evaluated how well-calibrated the confidence intervals were by calculating coverage, or the proportion of genes for which the true gene prevalence value *π_j_* fell within the confidence interval of the estimate *π_j_*. Both methodologies exhibited comparable coverage greater than or equal to the confidence level (Supplementary Figure 3C), a trend that held true across all gene prevalence categories at the 95% confidence level, with an overall coverage of almost exactly 0.95 for the MLE confidence intervals (Supplementary Figure 3D). Together with the evaluation of estimator accuracy (Figure 2), these results indicated that the MLE approach provided the most accurate estimates of gene prevalence when the dataset includes a significant number of incomplete genomes.

### Robustness to Model Mis-specification

We recognize that real presence/absence data will deviate from the described model due to factors such as inaccurate estimates of genome completeness and potential sequencing biases [28]. We therefore evaluated the effect of these potential violations of model assumptions on MLE gene prevalence estimates. We used simulated data to quantify the effect of violations in terms of overall error (MSE) and change in confidence interval coverage at the 95% confidence level.

#### Inaccurate Completeness Values

Genome completeness is an estimated value. While there are different approaches to estimating genome completeness [25, 29], most methods take annotated genes as inputs, particularly single copy core genes (SCGs), which are expected to be found in all representatives of a given taxonomy. In *Prochlorococcus* there are currently 1,128 SCGs listed in the latest version of the benchmarking sets of universal single-copy orthologs database (BUSCO v6.1.0) [29]. We simulated genome completeness values with inaccuracy levels that resulted from using different numbers of SCGs to estimate genome completeness. At 100 SCGs or more, the MSE between the true gene prevalence values and the estimates was nearly identical to the lowest error observed at 3,000 SCGs (Figure 3A). Similarly, the confidence intervals in simulations with 100 SCGs or more exhibited nominal coverage (Figure 3B). This indicates that under the MLE approach, inaccuracies in the genome completeness estimate are unlikely to significantly affect gene prevalence estimates when at least 100 SCGs are used to estimate genome completeness, a threshold exceeded in our *Prochlorococcus* dataset.

**Figure 3.**
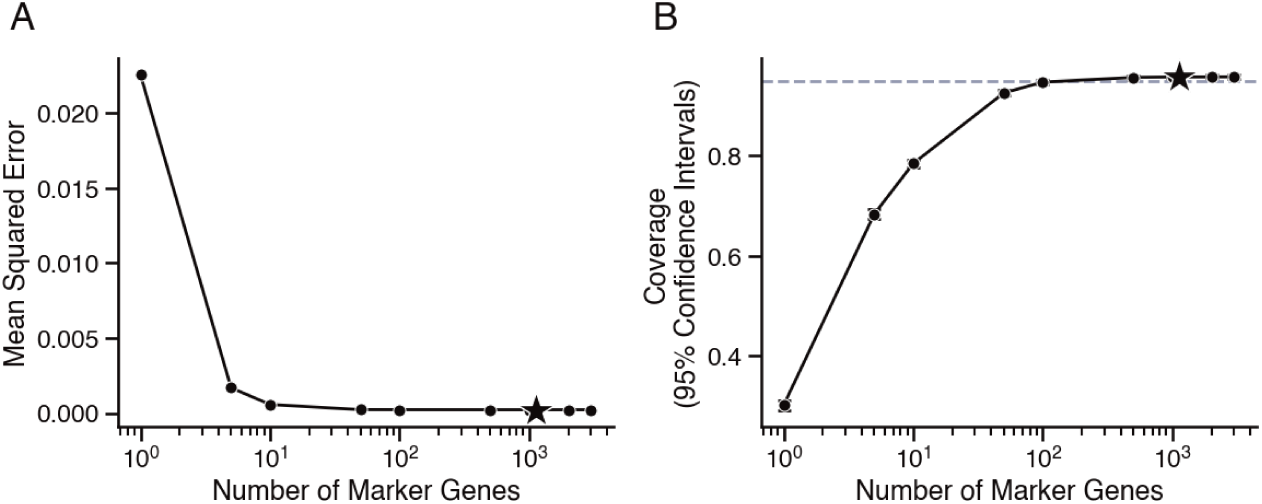
Simulation experiments show MLE method is robust to inaccurate genome completeness estimates. A) Mean Squared Error (MSE) and B) coverage (proportion of genes for which the true prevalence value falls within the 95% confidence interval of the estimate) from model estimates fit with simulated inaccurate genome completeness values. The degree of inaccuracy is correlated with the number of marker single-copy core genes (SCGs) used to estimate completeness, with more SCGs resulting in a more accurate completeness estimate. Star marks the number of SCGs available for *Prochlorococcus* in BUSCO v6.1.0.

#### Unequal Probability of Gene Detection

Our model supposes that every gene has an equal and independent chance of being observed. In reality, however, factors such as sequencing artifacts and multicopy genes may cause different gene families to be differentially detected. We evaluated the effect of differential detection on the performance of our model by simulating sets of incompletely sequenced genomes in which some genes were systematically more likely to be observed than others (see Methods). The model was robust to low levels of variation in gene detection probability. As variation increased, overall model performance decreased, with estimate error increasing and confidence interval coverage dropping below the nominal 95% confidence level threshold (Supplementary Figure 4).

Given these findings, we quantified the effect on our model of multicopy genes, which likely result in variation in gene detection probabilities. Genes found in more than one copy per genome are expected to be detected in incomplete genomes more frequently than single copy genes. We tested this effect with simulations in which the copy number of each gene family was systematically varied, testing scenarios ranging from the average gene family having a copy number of 1 (i.e. all genes are single copy) to the average gene family having 11 copies per genome (Figure 4A). For comparison, the average *Prochlorococcus* gene family has a copy number of 1.023 in our dataset, an underestimate of the true value due to genome incompleteness. As expected, gene prevalence estimates diverged from ground truth as average gene copy number increased (Figure 4B, C), and confidence interval coverage dropped (Figure 4D). However, at a mean copy number of 1.023, coverage remained above the nominal threshold and MSE was comparable to the single copy gene scenario, indicating that in the case of *Prochlorococcus*, gene copy number is unlikely to substantially impact model performance across the pangenome as a whole. That being said, we found 369 *Prochlorococcus* gene families with a mean copy number of 2 or greater, and a maximum mean copy number of 13. In these cases, our simulation experiments revealed elevated error and a breakdown in confidence interval coverage, particularly for genes of intermediate prevalence (Supplementary Figure 5). We therefore aimed to update the model to account for bias introduced by multicopy genes.

**Figure 4.**
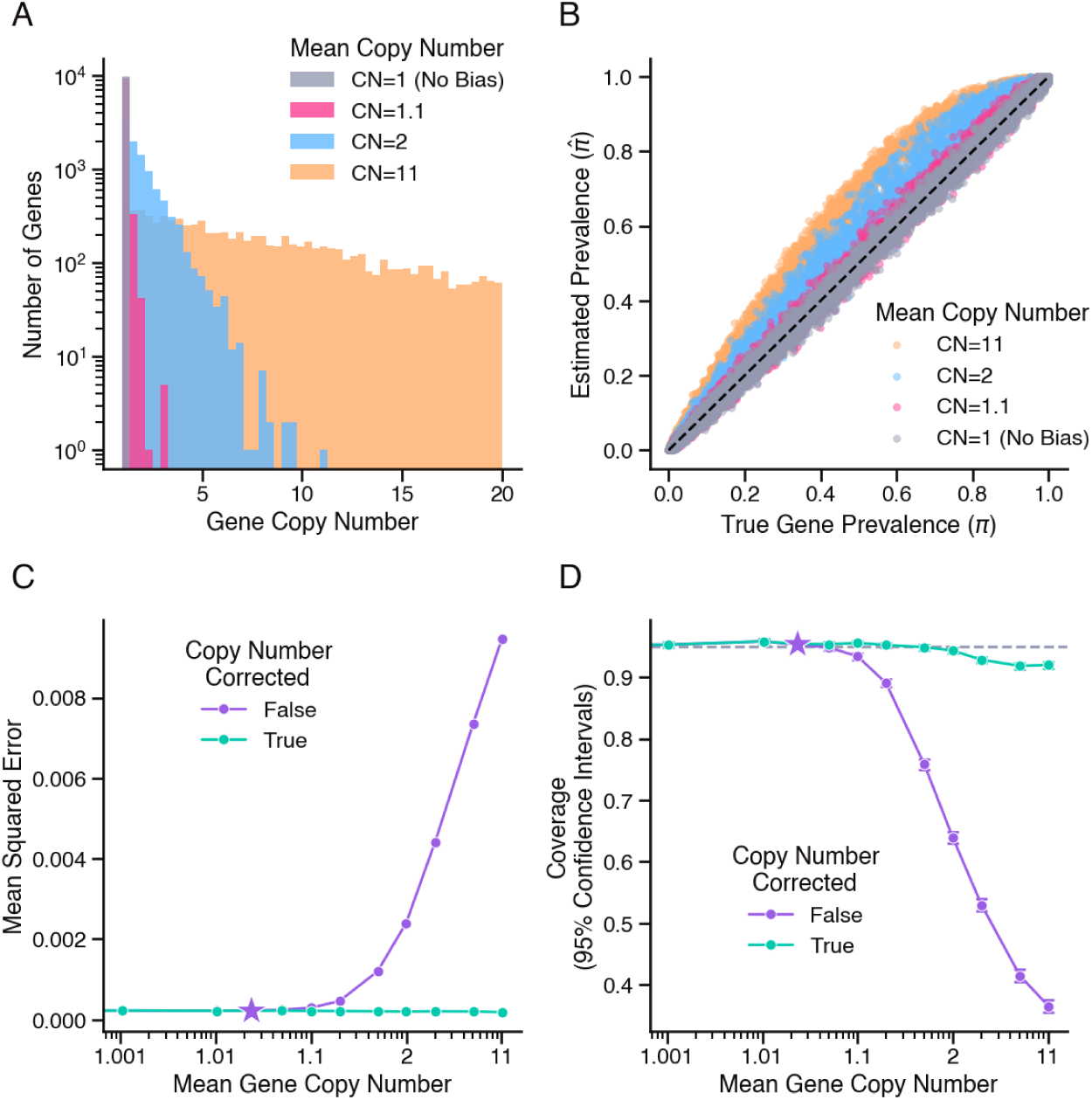
Simulation experiments show MLE method is robust to biased gene detection resulting from multicopy genes. A) Distribution of true simulated gene copy numbers (i.e. 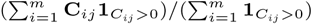). Color indicates overall mean gene copy number (CN) across simulated pangenome as a whole. B) Estimated versus true gene prevalence from models fit to presence/absence data generated from simulated pangenomes with multicopy genes. C) Mean squared error (MSE) and D) coverage (proportion of genes for which the true prevalence value falls within the 95% confidence interval of the estimate) from model estimates fit to simulated pangenomes with increasing overall mean gene copy number. Line color represents model estimation with and without adjusting for copy number. Bars indicate 2× standard error. Star marks overall mean gene copy number in the real dataset of available *Prochlorococcus* genomes (1.023).

To further improve model robustness and expand applicability to gene families with mean copy numbers greater than 1, we modified our model of the observation matrix **Z**. In the updated version, each entry **Z***_ij_* is the result of **C***_ij_* independent Bernoulli trials, where **C***_ij_* is the copy number of gene *j* in genome *i* (see Methods). We updated the MLE likelihood function with this copy number correction and evaluated the change in performance with simulated data, using the average observed copy number of each gene family as an estimate of the true underlying **C***_ij_* parameter. The updated model successfully controlled error, even at the highest average copy number tested (Figure 4C), and maintained nominal confidence interval coverage for simulated pangenomes with an average gene copy number of 1.5 or less (Figure 4D). In the simulation with a mean copy number of 1.023, the MSE improved from 2.25 × 10^−4^ to 2.19 × 10^−4^, driven by improved accuracy in estimating the prevalence of gene families with elevated mean copy numbers (Supplementary Figure 5). This indicates that even for organisms like *Prochlorococcus* with relatively low levels of multicopy genes [30], the copy number corrected model offers more accurate prevalence estimates for those gene families present in multiple copies per genome.

#### Evaluation with Real Data

We next evaluated model performance with real data using two independent samples of genomes from our compiled *Prochlorococcus* dataset: one drawn exclusively from the completely sequenced closed genomes (*m* = 50), and the second drawn from the incomplete genomes (*m* = 50). We applied maximum likelihood estimation to each sample independently. The resulting gene prevalence estimates for these two data subsets were generally comparable (Supplementary Figure 6A). Specifically, the gene prevalence estimate based on the complete genomes exceeded the estimate from the incomplete genomes for 48.9% of gene families, fell below for 40.5% of gene families, and was exactly equal for 10.7% of gene families (Supplementary Figure 6B). The gene families for which the two estimates were equal were all core genes with predicted prevalence values of 1. Furthermore, at a 95% confidence level, the gene prevalence estimate from one sample fell within the confidence interval of the estimate from the other sample for 96.6% of gene families, indicating that confidence intervals remain well-calibrated in our dataset. These results suggest that model assumptions are well-suited to the data, and that the methodology generalizes well to patterns of missingness encountered in real data.

### Improvement Over Baseline

Use of both complete and incomplete genomes allows for a more extensive sampling of the set of genes that comprise a pangenome. We performed pangenome analysis on the full set of 1,099 complete and incomplete *Prochlorococcus* genomes, revealing a total of 52,500 unique gene families, including 40,516 gene families detected only in the incomplete genomes (Supplementary File 2). Historically this latter group has been excluded from gene prevalence estimation, and thus our methodology extended gene prevalence quantification to over four times the number of genes that are quantifiable when limited to complete genomes (Figure 5A). Furthermore, the additional genomic information enabled higher confidence estimates for all genes in the pangenome, with the median 95% confidence interval width shrinking from 1.23 × 10^−1^ when limited to complete genomes to 6.81 × 10^−3^ when using the full set of available *Prochlorococcus* genomes (Figure 5B). These trends underscore that our methodology enables a broader set of genomic data to be incorporated into pangenome analyses, resulting in a more complete and higher confidence survey of gene prevalence across the pangenome.

**Figure 5.**
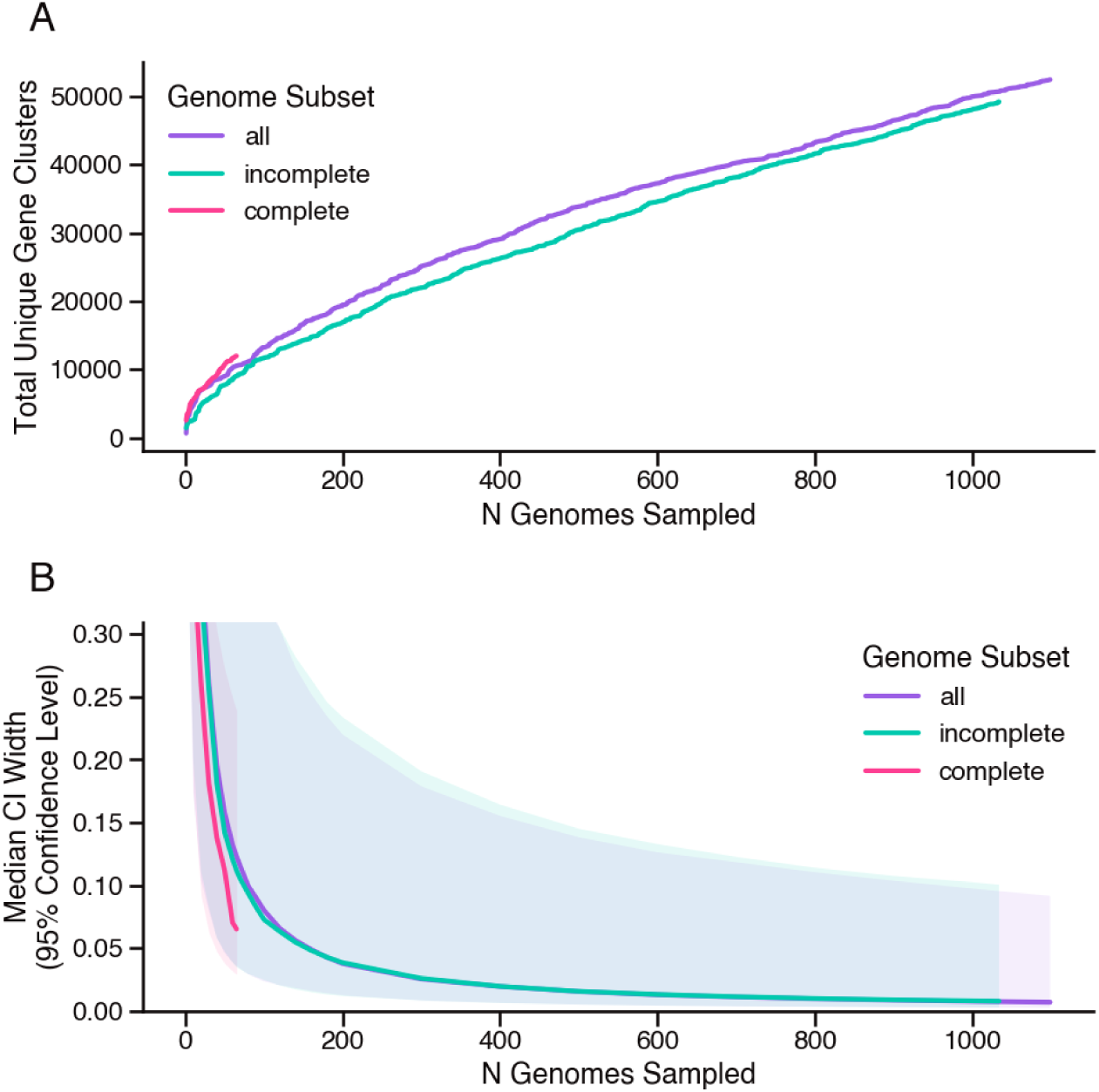
Methodology enables higher confidence gene prevalence estimates for a larger fraction of the *Prochlorococcus* pangenome. A) Saturation curve illustrating the total number of unique genes present in subsets of the complete (pink), incomplete (green) and combined (purple) *Prochlorococcus* genomes. B) Median width of 95% confidence intervals (CIs) for gene prevalence estimates from MLE model fit to presence/absence data derived from indicated subsets of *Prochlorococcus* genomes. Bands indicate total range of CI widths.

### Combining Subpopulation Estimates

The *Prochlorococcus* genus is subdivided into ecologically differentiated subpopulations, including the high-light clades (e.g. HLI and HLII) better adapted to surface waters, and the low-light clades (e.g. LLI and LLVII) more frequently found lower in the water column [4, 31]. The global *Prochlorococcus* population is comprised of a mix of these different clades (Supplementary File 1). Our methodology assumes that the input genomes are a representative sample of the population of interest, implying proportional representation of the different *Prochlorococcus* clades. Although single-cell genome sequencing can reduce some sampling bias, other biases remain from factors such as culture selection and non-random sampling locations, making it unlikely that the clade composition of the genome dataset is proportional to the global *Prochlorococcus* population. We therefore evaluated how gene prevalence varied between the different clade subpopulations of *Prochlorococcus* in order to determine whether to treat our set of genomes as representative of one homogeneous population, or to consider clade subpopulation individually.

We generated two subsamples of genomes, one drawn exclusively from genomes belonging to the HLII clade of *Prochlorococcus* (*m* = 50), and the other from the LLI clade (*m* = 50). We applied our maximum likelihood estimator to each sample independently, and compared gene prevalence estimates between the two datasets (Supplementary Figure 7A). The resulting distribution revealed three distinct subsets of gene families: a set found only in HLII genomes, a second set found only in LLI genomes, and a third set found at high prevalence in both sets of genomes. In contrast, two independent samples both drawn from the HLII genomes resulted in gene prevalence values that were generally consistent between the two samples (Supplementary Figure 7B). This indicates that while gene prevalence values are generally homogeneous within subpopulations of the same *Prochlorococcus* clade, gene content diverges between different clades, necessitating a separate consideration of clade subpopulations when estimating gene prevalence.

Estimates of gene prevalence in heterogeneous populations are key to identifying core genes across, for example, all *Prochlorococcus* clades. Therefore we developed a methodology for combining prevalence estimates of subpopulations, given the relative abundance of each subpopulation in the target population (see Methods). In brief, overall prevalence can be calculated as a weighted average of the per-subpopulation prevalence estimates. To construct confidence intervals for the overall population, in each subpopulation we approximated the standard deviation of the prevalence estimate using its confidence interval and propagated these uncertainties through the weighted average. Because the subpopulation-specific estimates are independent, their estimation errors can partially cancel, potentially yielding narrower confidence intervals for the combined population than for individual subpopulations. This approach treats the relative abundances of the subpopulations as fixed and does not account for uncertainty in their estimation. We next applied this methodology to our dataset of genomes to estimate the global prevalence of every detected gene in the *Prochlorococcus* pangenome.

### Estimating Global *Prochlorococcus* Gene Prevalence

Estimates of global gene prevalence values for the *Prochlorococcus* pangenome require knowledge of the relative abundance of each clade in the global *Prochlorococcus* population. To approximate these relative abundances, we leveraged a global dataset of 16S rRNA amplicon sequence variants (ASVs) [32]. We identified all the ASVs that originated from *Prochlorococcus* cells and annotated each with clade of origin, using as our reference the compiled dataset of *Prochlorococcus* genomes (Supplementary File 3). Samples were then grouped into nine bins based on the Longhurst biome [33] and depth of the sampling location. The relative abundance of each clade was then aggregated to obtain an estimate of the mean clade composition of the *Prochlorococcus* population within each biome–depth bin (Supplementary Figure 8, Supplementary File 4). We compared these bins against an unconstrained data-driven clustering approach and found that while the latter approach produced more homogeneous clusters of ASV profiles, the total variance explained was comparable between the two approaches (Supplementary Figure 9), validating that the binning approach captures much of the reproducible variation in the global *Prochlorococcus* population structure. We then used an annual climatological model of average *Prochlorococcus* cell counts to estimate the relative sizes of the subpopulations in each biome–depth bin (Supplementary Figure 8) and aggregated across bins to produce a single estimate for the clade composition of the global *Prochlorococcus* population (Figure 6). This estimate revealed that the global population is dominated by the HLII (43.8%), LLI (18.2%), and HLI (15.9%) clades, whereas none of the other clades were found to make up more than 5% of the global population. A total of 12.8% of the global population corresponded to ASVs that could not be unambiguously annotated with a single clade, suggesting that these estimates represent a lower bound on the true relative abundance values, and that a small subset of the *Prochlorococcus* population may belong to clades that are yet to be characterized.

**Figure 6.**
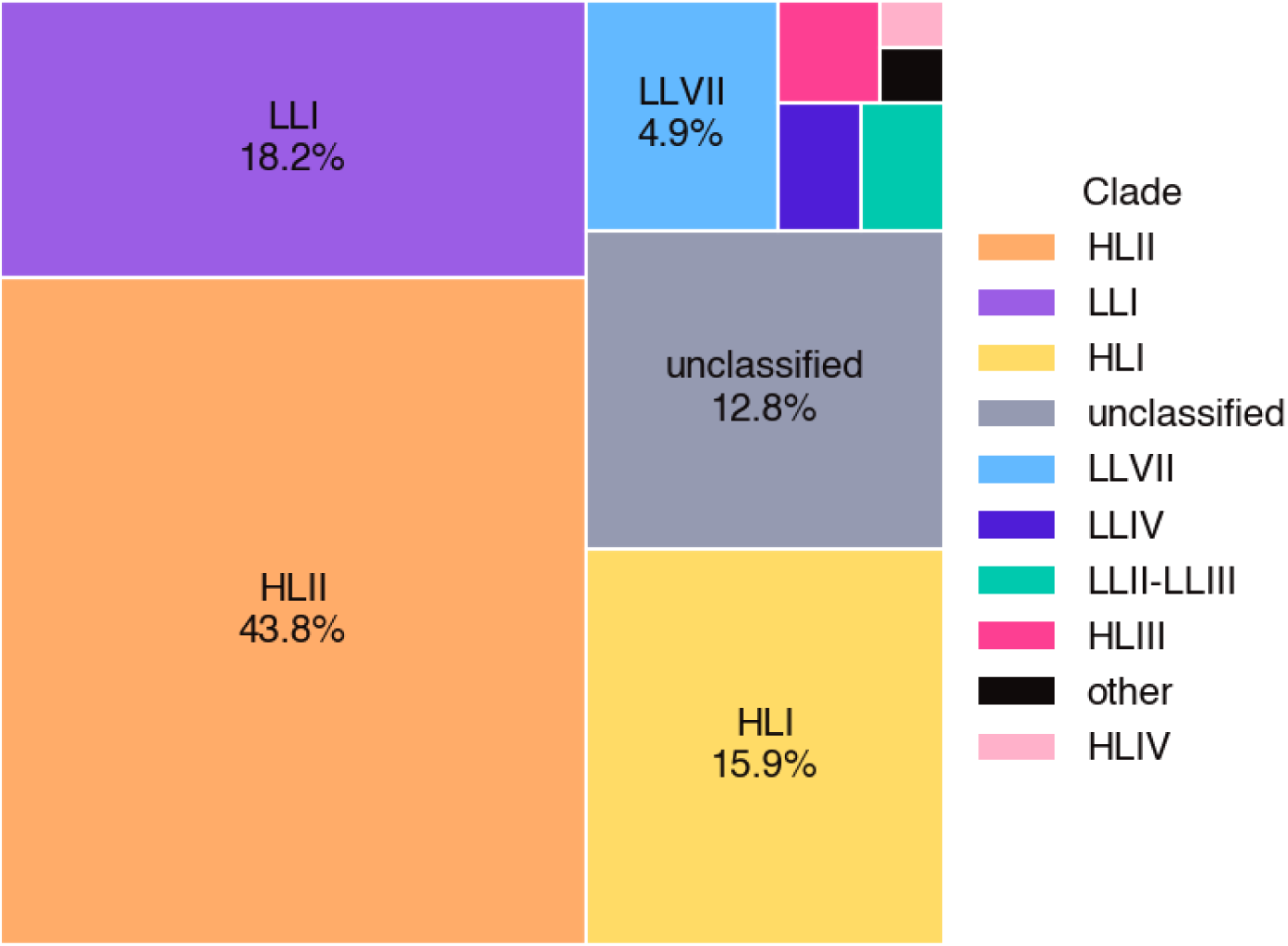
High light clades constitute a majority of the global *Prochlorococcus* population. Color indicates clade and rectangle area indicates relative abundance of clade as estimated from global dataset of 16S rRNA amplicon sequence variants [32].

We generated estimates for the global prevalence of each gene family in the *Prochlorococcus* pangenome by aggregating estimates from each clade subpopulation. First, the maximum likelihood estimator was independently applied to the subset of genomes belonging to each clade (Supplementary File 5). Then, the resulting clade-specific gene prevalence estimates were aggregated, weighting by the estimated relative abundance values of each clade to produce the global gene prevalence estimates (Figure 7, Supplementary File 6). Surprisingly, only 27 gene families were estimated to have a global prevalence of exactly 1. However this point estimate does not account for routine sources of error in sequencing, gene calling, and other aspects of sampling, so the confidence intervals provided by our methodology were leveraged to incorporate uncertainty. A gene family was defined as ‘core’ when its 95% confidence interval included a prevalence of 1. By this definition we found that the core of the *Prochlorococcus* pangenome consists of 714 gene families. Consistent with *Prochlorococcus* having an open pangenome, the vast majority of gene families were cloud genes (prevalence ≤ 0.15; n=49,828) and 1,708 gene families fell into the shell category (0.15 *<* prevalence *<* 0.9). The extended core (prevalence ≥ 0.9) consisted of a total of 964 gene families. Within clade subpopulations, the size of the core ranged from 1,113 to 2,604 gene families, revealing that the number of core genes within each clade is larger than the number of genes core to the *Prochlorococcus* population overall (Supplementary Figure 10). The total number of cloud genes in each clade was larger in clades for which there were more available genomes, underscoring that the vast majority of protein-coding genetic material in the *Prochlorococcus* pangenome consists of low-prevalence flexible genes. Altogether, these results show that the global *Prochlorococcus* pangenome consists of a relatively small set of broadly distributed genes and a much larger pool of genes with prevalences that vary between clades and across the global population.

**Figure 7.**
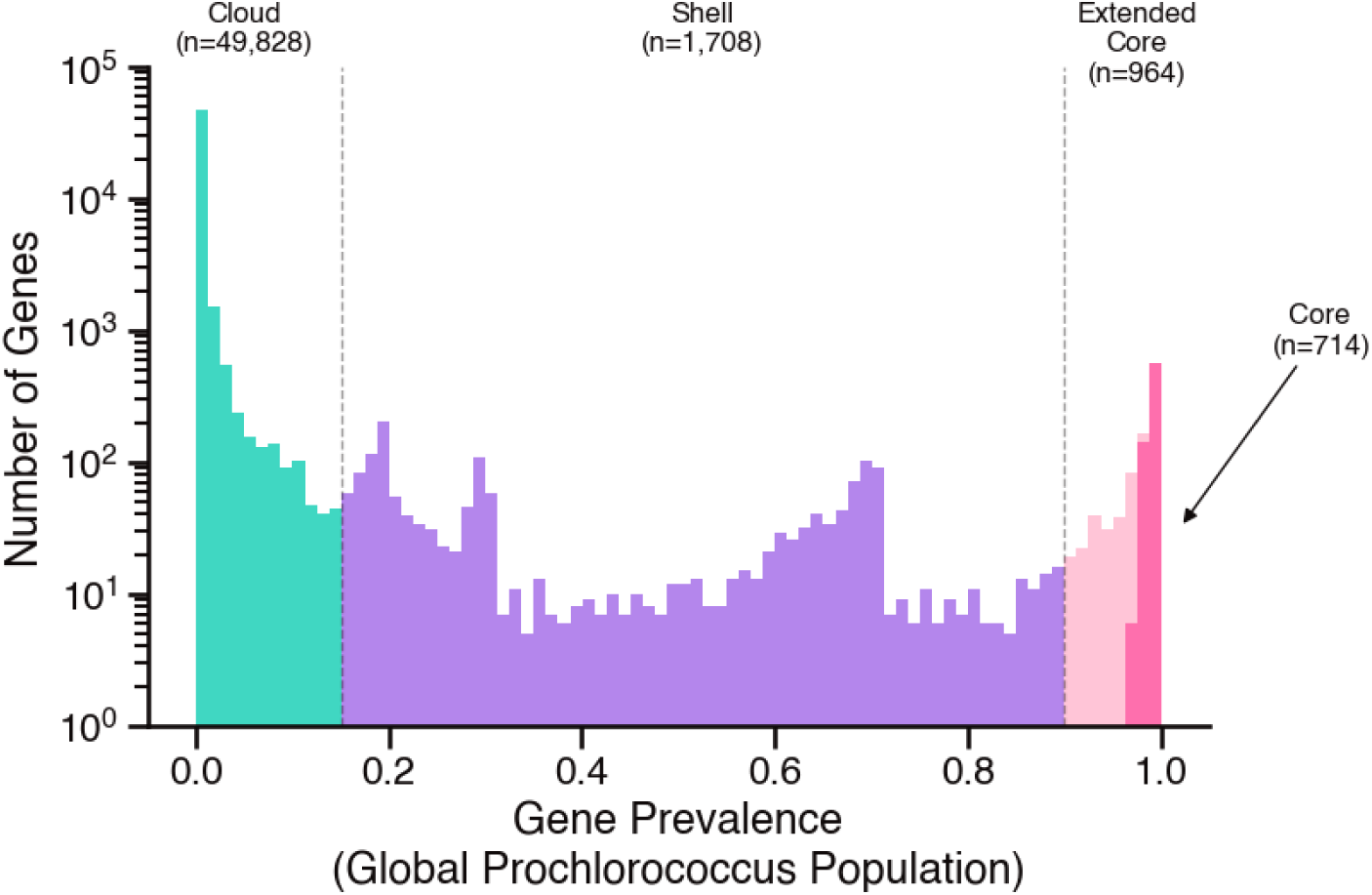
Global *Prochlorococcus* pangenome predominantly consists of rare flexible cloud genes, with a core of 714 gene families that are likely universal. Estimates of gene prevalence in the global *Prochlorococcus* population, colored by pangenome category.

## Discussion

In this study we developed a statistical framework for estimating the prevalence of gene orthologs within a population of interest, given a sample of complete and potentially incomplete genomes. Previous approaches to pangenome analysis commonly either did not incorporate genome completeness information into gene prevalence calculations [16, 34] or else limited analysis to genomes that passed a completeness threshold [15, 17]. The first approach results in a systematic under-estimation of gene prevalence [18] and the second limits the scope and accuracy of the resulting analysis (Figure 5). Our methodology improves upon such approaches by modeling the observed presence/absence pattern of each gene family as a Bernoulli process governed by the completeness level of each genome in the dataset and a true underlying gene prevalence value, the latter parameter being the estimation target. There are existing methods that account for genome completeness [24, 23], including CELEBRIMBOR, which uses the same fundamental statistical model as described here [20]. However, these methods aim to classify genes into core and accessory categories (sometimes further distinguishing between shell and cloud genes), rather than quantifying the prevalence and associated uncertainty of each gene family. By incorporating genomes across a range of completeness levels, our approach makes fuller use of available genomic data and provides quantitative estimates of gene prevalence and associated uncertainty.

We applied our methodology to a dataset of 1,099 *Prochlorococcus* genomes, of which 94% were incomplete isolate genomes and SAGs (Supplementary Figure 1). Our analysis produced estimates of the global prevalence of 52,500 unique gene families (Supplementary File 6), the broadest investigation of the *Prochlorococcus* pangenome reported to date. The overwhelming majority of gene families (n=49,828) were low prevalence cloud genes, consistent with the characterization of the *Prochlorococcus* pangenome as ‘open’ compared to those of other microbial taxa [3]. Additionally, we found that every genome in the compiled dataset carried novel gene families not yet detected in any other genome (Figure 5A). This observation suggests that many *Prochlorococcus* genes, perhaps a majority, are yet to be sequenced. The large tail of rare cloud genes comprises the leftmost peak of the U-shaped gene prevalence distribution typical of prokaryotic pangenomes [26, 27], while a smaller peak near 1 corresponds to the universal or near-universal core genes (Figure 7). Previous approaches to classifying core genes aimed to mitigate errors in sampling, sequencing, and gene calling by fine-tuning thresholds to be more lenient than absolute universality [13, 19, 20]. The confidence intervals produced by our method allow for a more statistically grounded treatment of uncertainty in this delineation. We defined core genes as those for which the 95% confidence interval included a prevalence of 1, resulting in a core of 714 genes. This number is consistent with previous investigations of the *Prochlorococcus* pangenome [14], including the most recent analysis with 31 isolate genomes that identified a core of 766 genes [15].

Between the peaks at 0 and 1 corresponding to cloud and core genes, the *Prochlorococcus* gene prevalence distribution also had several smaller intervening peaks within the range of the shell genes (Figure 7). A previous study cited this pattern as evidence that the *Prochlorococcus* taxon should be divided into two or more species, each with internally homogeneous gene content [35]. This view is supported by our finding that gene prevalence estimates are consistent between subsamples of the same clade whereas they diverge in subsamples of different clades (Supplementary Figure 7). This pattern is likely a result of established niche differentiation between different clades of *Prochlorococcus* [36, 4], which was reiterated in the relative abundance patterns in our global analysis of amplicon sequence variants [32]: the HLII clade dominates in the Trades biome near the equator, HLI is relatively more abundant in the colder waters of the Westerlies biome, and the low light clades are relatively more abundant lower in the water column (Supplementary Figure 8, Supplementary File 4). Because of the variation in gene content between clades, the prevalence of a gene in the overall *Prochlorococcus*population depends in part on the relative abundance of clade subpopulations. This informed our approach to estimate gene prevalence independently within each clade (Supplementary File 5), before combining these estimates for an overall estimate of gene prevalence in the global *Prochlorococcus* population (Figure 7, Supplementary File 6). More broadly, these findings illustrate how the interpretation of a pangenome depends on the population over which it is defined. Recent work has argued for considering constrained local populations in pangenome analysis, as the flexible gene pool shared among coexisting cells reflects genomic variation within a common ecological and evolutionary context [7]. From this perspective, clade-specific and global *Prochlorococcus* pangenomes have distinct biological interpretations: the former characterize gene distributions within more ecologically and evolutionarily cohesive populations, whereas the latter characterize gene distributions across the diversity of the taxon as a whole.

The framework developed here provides a foundation for further improvements in gene prevalence estimation and opens the door for new approaches to investigating microbial genomic diversity. Incomplete genomes represent an important resource in these efforts, as they enable genomic sampling at greater scale and expand the diversity of organisms and environments represented. The presented methodology enables incomplete genomes to be incorporated into estimating gene prevalence, although opportunities remain to improve accuracy and applicability. Our experiments with simulated data demonstrated that explicitly correcting for multiple copies of genes within genomes resulted in more accurate estimates (Figure 4), suggesting that incorporating other available information could further improve estimation. For instance, Mattick et al. recently proposed a phylogenetic occupancy model that incorporates evolutionary relationships among genomes when inferring gene presence from incomplete data [37]. Estimates might similarly be improved by incorporating information about gene synteny, as PPanGGOLiN does using a graph-based framework [24], as well as the evolutionary histories of individual gene families or patterns of gene co-occurrence. In combination with continued improvements in sequencing technology, these and other methodological advances will enable innovative applications of gene prevalence quantification to questions of microbial ecology and evolution. For instance, in addition to SAGs and MAGs, advances in long-read metagenomics are being deployed to assess flexible gene content directly within natural populations [38, 39]. This approach could be applied to time-series data, in which case the methodological framework outlined here could be leveraged to track changes in gene prevalence within a local microbiome, providing a quantitative approach for investigating the ecological and evolutionary dynamics that shape microbial gene pools through time. Altogether this work highlights the value of incorporating incomplete genomes into quantitative pangenome analysis and the broader potential of gene prevalence as a measure of microbial genomic diversity.

## Methods

### Model

We developed a statistical model to formalize the conceptual model that each observation of a gene family in a genome is the result of 1) the gene being truly present in that genome and 2) a successful detection event, including sequencing, gene annotation, etc. Let **X** ∈ {0, 1}*^m×n^* represent the observed gene family presence/absence data, where *m* is the number of genomes and *n* the number of gene families. We index genomes using *i* = 1*, …, m* and gene families using *j* = 1*, …, n*. Each presence/absence observation **X***_ij_* = 1 if gene family *j* was observed in genome *i*, and **X***_ij_* = 0 otherwise. We modeled this matrix as an entry-wise (i.e. Hadamard) product of a true presence/absence matrix **Y** ∈ {0, 1}*^m×n^* and an observation matrix **Z** ∈ {0, 1}*^m×n^*. In other words, for each genome *i* and each gene family *j*, **Y***_ij_* indicates whether the gene is truly present and **Z***_ij_* whether there was a successful detection event, thus yielding:

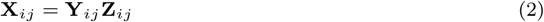

Our quantity of interest is the true prevalence of each gene family. We denote the true gene prevalence of gene family *j* as *π_j_*, defined as the probability that the gene is present in a genome randomly sampled from the population. We modeled true gene presence/absence as **Y***_ij_* ∼ Bernoulli(*π_j_*) for all genes *j* and genomes *i* independently across genes and genomes. Similarly, we modeled each observation event as a second set of independent Bernoulli random variables with **Z***_ij_* ∼ Bernoulli(*ζ_ij_*), where *ζ_ij_* is the probability that the status of gene family *j* was observed in genome *i*, or equivalently, *ζ_ij_* = *P* (**Z***_ij_* = 1). We can relate *ζ_ij_* to the completeness of genome *i*, denoted as *λ_i_*, with one of two simplifying assumptions. The simplest assumption is that *ζ_ij_* depends only on genome *i*, so that every gene in genome *i* has the same chance *λ_i_* of being observed, i.e., assumption:

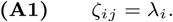

In other words, in a genome estimated to be 80% complete, **(A1)** posits that every truly present gene has an 80% chance of being observed. This also means that **Y***_ij_* is independent of **Z***_ij_*. This assumption can be violated if the genomes have multiple copies of genes. To account for gene copy number, let **C***_ij_* denote the copy number of gene *j* in genome *i*, and let **Y***_ij_* = **1_C_***_ij_ ≥*1. In other words, **Y***_ij_* = 1 if gene *j* is found in genome *i* with a copy number of 1 or greater, and otherwise **Y***_ij_* = 0. If we now assume that each copy of gene *j* in genome *i* is observed or missed independently with the same underlying probability of *λ_i_*, then **Z***_ij_* = 0 if **C***_ij_* independent Bernoulli(*λ_i_*) trials all return 0, or equivalently:

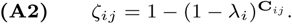

Note that the assumption **(A1)** is a special case of assumption **(A2)**; if **C***_ij_* = 1 for all genomes *i* and genes *j* then **(A2)** implies **(A1)**. Also note that in this case **Z***_ij_* is not a priori independent of **Y***_ij_*; both depend on the unobserved counts **C***_ij_*.

Further, we assumed that we have access to the completeness parameter *λ_i_* for each genome, or at least a sensible estimate of this value. We did not assume knowledge of the true gene counts for each genome, but instead assumed that we have access to an estimate of the average per-gene copy number such as 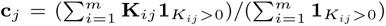 where **K***_ij_* are the observed gene counts, with distributions given by **K***_ij_* | **C***_ij_* ∼ Binomial(**C***_ij_, λ_i_*). Thus we will not know *ζ_ij_* unless we make assumption **(A1)**.

### Estimators

We considered four approaches to estimating gene prevalence. These differ primarily in how they account for incomplete genomes: the naive estimator ignores incompleteness, the subset estimator excludes incomplete genomes, the unbiased estimator weights observations according to genome completeness, and the maximum-likelihood estimator explicitly models the observation process.

#### Naive Estimator

We formulated the naive estimator to ignore genome incompleteness and calculate the gene prevalence estimates as the column means of the observed presence/absence data. Mathematically this is equivalent to:

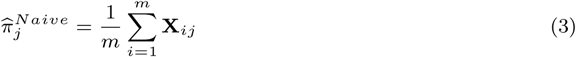

By ignoring the missingness in the incomplete genomes, we assume that the naive estimator will underestimate true gene prevalence.

#### Subset Estimator

Alternately, we formulated the subset estimator to ignore the data in the incomplete genomes and focus only on the complete genomes. We express this estimator as:

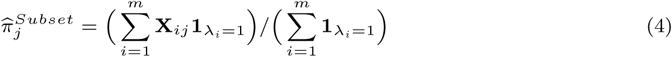

This approach avoids the downward bias of the naive estimator but has larger variance because it ignores the information in the incomplete genomes.

#### Unbiased Estimator

Rather than discarding incomplete genomes, we can instead compensate for the lower probability of detecting genes in these genomes as compared to more complete genomes. In this setting, an observed presence in genome *i* with completeness *λ_i_* is therefore weighted by 1*/λ_i_*. This approach is similar to inverse probability weighting techniques used elsewhere to account for missing data [40], and we formulate it here as follows:

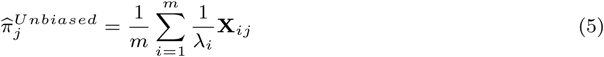

To see that this estimator is unbiased, note that under assumption **(A1)**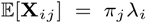 and so 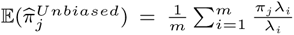. However, it’s worth noting that this estimator can be biased when *ζ_ij_* ≠ *λ_i_* for some *i, j*. Additionally, when the completeness values *λ_i_* are small, this estimator can produce values greater than 1. This can be corrected by clipping the estimator:

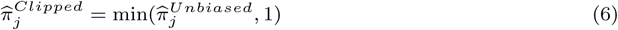

This clipped version of the unbiased estimator will be closer to the true prevalence *π_j_* for every possibly sampled data **X**, so this clipping procedure reduces the mean squared error through a targeted bias-variance tradeoff. That is to say we introduce a small amount of bias at the high end to get an outsized reduction in the variance caused by estimates that are much greater than 1.

#### Maximum Likelihood Estimation

As an alternative to correcting individual observations, we can estimate gene prevalence by identifying the value of *π_j_* under which the observed pattern of gene presences and absences is most likely, given the completeness of the sampled genomes. This gives the maximum likelihood estimator (MLE). Under our model, the probability of observing gene *j* in genome *i* is *π_j_ζ_ij_*, while the probability of not observing it is 1 − *π_j_ζ_ij_*. Assuming independence across genomes, the likelihood of the observed data for gene *j* can therefore be written as:

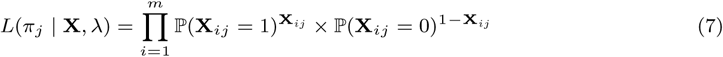

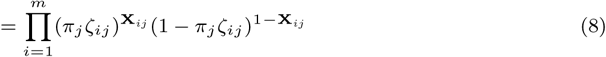

For numerical optimization, it is more convenient to maximize the logarithm of the likelihood. Because the logarithm is monotonic, this produces exactly the same estimate of *π_j_*, while converting products into sums:

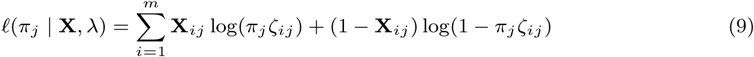

The maximum likelihood estimator is then defined by the maximizer:

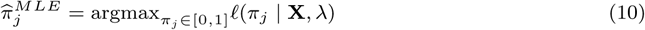

We chose to find the maximum by considering the derivative in *π_j_* and solving where this derivative equals 0:

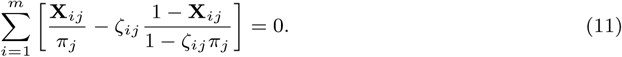

Note that this also requires consideration of the values at the boundaries *π_j_* = 0 and *π_j_* = 1. Because this is a constrained optimization procedure we will never have to clip this estimator. The values used for *ζ_ij_* will depend on the assumption **(A1)** or **(A2)**. In the case of assumption **(A1)** the MLE would use *ζ_ij_* = *λ_i_*, and under **(A2)** we would use *ζ_ij_* = 1 − (1 − *λ_i_*)**^c^***j*. In the second case we are using the estimated average copy number per gene rather than the true gene counts **C***_ij_*, which we are unable to observe whenever there are incomplete genomes in the sample. Note also that because we assumed independence of every *Y_ij_* the MLE of *π_j_* only depends on the *j^th^* column of **X**; the other gene families have no bearing on this estimate. Consequently, this estimator is automatically robust to deviations from the independence assumption where **Y***_ij_*_1_ can correlate with Y*_ij_*_2_ (i.e., if the genes can co-occur), however, the estimator would no longer be the MLE given the data **X** and would instead be interpreted as the MLE given the data in the *j^th^* column of **X**.

### Confidence Intervals

In addition to estimating gene prevalence, we sought to quantify the uncertainty associated with each estimate. We used two approaches to construct confidence intervals, corresponding to the different estimators described above. For the subset and unbiased estimators, we constructed intervals from the estimated variance of the prevalence estimator, while for the MLE we used likelihood ratio intervals.

#### Variance Based Intervals

For the subset and unbiased estimators, uncertainty in the prevalence estimate can be quantified from the variance expected under our model. Under assumption (A1), the variance of the unbiased estimator is given by:

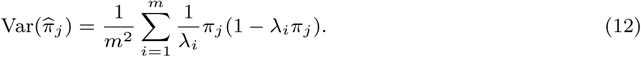

Because the true prevalence *π_j_* is unknown, we estimate this variance by substituting the corresponding prevalence estimate 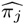:

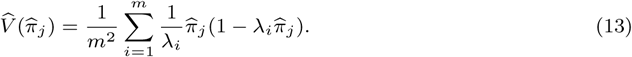

We then approximated the distribution of *π̂_j_* as a Gaussian centered at *π_j_* with variance Var(*π̂_j_*). Specifically, we constructed confidence intervals at the (1 − *α*)-confidence level as follows:

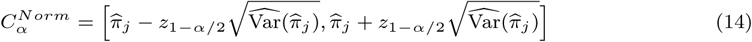

where *z*_1_*_−α/_*_2_ is the 1 − *α/*2 quantile of a standard normal distribution (e.g., *z*_0.975_ ≈ 1.96 for a 95% confidence interval).

#### Likelihood Ratio Intervals

The MLE identifies the single prevalence value that maximizes the likelihood of the observed data, but nearby prevalence values may provide nearly as much support for the observations. We therefore constructed confidence intervals by identifying the range of prevalence values for which the log-likelihood remained sufficiently close to its maximum. Specifically, we included values of *π_j_* satisfying:

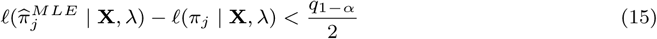

where *q*_1_*_−α_* is the 1 − *α* quantile of a 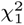 distribution.

Classical likelihood-ratio confidence intervals rely on regularity conditions that are not strictly satisfied by our model. In particular, genomes can have different completeness values *λ_i_*, such that observations are independent but not identically distributed, and the true prevalence *π_j_* may lie on the boundary of its parameter space at 0 or 1. We therefore evaluated the empirical coverage of these intervals using simulations (see Results). In practice we do not see that these intervals exhibit any further type 1 error. For interior parameter values the MLE still appears sufficiently approximately normal to get valid confidence intervals in all of our simulations, and on the boundary the test fails safely where we see conservative confidence intervals.

#### Combining Confidence Intervals

In some applications, we may wish to estimate gene prevalence in a population composed of multiple subpopulations each with different gene prevalence values. We therefore considered how prevalence estimates and their associated confidence intervals can be combined across subpopulations according to their relative abundance. Let *k* = 1*, …, K* index *K* subpopulations, with relative abundances *p_k_*, where *p_k_* >0 and 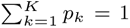. For gene *j*, let 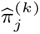 denote the prevalence estimate within subpopulation *k*. The combined prevalence estimate is simply the average of the subpopulation-specific estimates, weighted by relative abundance:

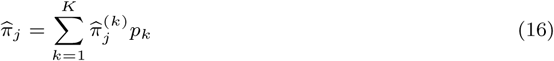

Combining the corresponding confidence intervals requires accounting for additional uncertainty in each of the subpopulation-specific estimates. Suppose that the prevalence estimate for gene *j* in each subpopulation *k* has a confidence interval 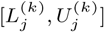. If all *K* subpopulation-specific intervals contain their respective true prevalence values, then the weighted average of their lower bounds and the weighted average of their upper bounds must contain the true combined prevalence. We can therefore construct a combined interval with the following bounds:

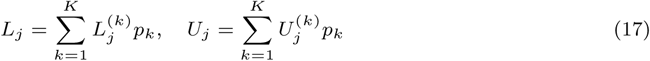

If each subpopulation-specific interval has coverage 1 − *α^′^* and the estimates are independent across subpopulations, then the probability that all *K* intervals simultaneously contain their respective true prevalence values is (1 − *α^′^*)*^K^*. To obtain a combined confidence interval with coverage of at least 1 − *α*, we therefore set

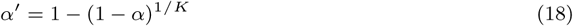

and construct each subpopulation-specific interval with confidence level 1 − *α^′^*. We refer to the resulting weighted interval [*L_j_, U_j_*] as the **guaranteed combined confidence interval**. For comparison, we also calculated an **uncorrected combined confidence interval** by applying the same weighted averaging of interval bounds without adjusting the confidence level of the subpopulation-specific intervals.

The guaranteed combined confidence interval can be conservative because it requires simultaneous coverage of the true prevalence in every subpopulation. We therefore also considered an approximate approach that propagates the estimated variances of the independent subpopulation-specific estimates through their weighted average. If 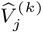 is the estimated variance of 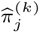, then the estimated variance of the combined prevalence is

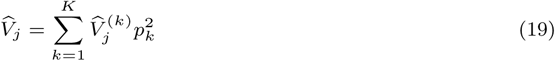

We then construct the combined confidence interval as

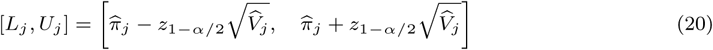

where *z*_1_*_−α/_*_2_ is again the corresponding quantile of the standard normal distribution. When 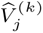 is not directly available (as with the likelihood ratio confidence intervals), we approximate it from the corresponding confidence interval as

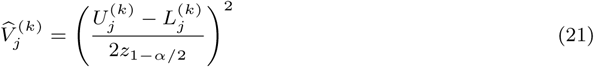

We refer to intervals constructed using this variance-based approach as the **approximate combined confidence interval**.

### Simulated Data

To evaluate the performance of estimation procedures under well-specified conditions and in cases where model assumptions were violated, we simulated presence/absence data for a pangenome of 1,000 genomes and 12,000 gene families. We drew a vector of 12,000 ground truth gene prevalence values *π* from a Beta distribution (*α* = 0.2*, β* = 0.5), multiplied this vector by 1.06, and clipped values to 0 and 1, to ensure a subset of core genes where *π_j_* = 1. We then used these true prevalence values to generate a ground truth gene presence/absence matrix via Bernoulli random draws, with each Bernoulli random variable parameterized by the corresponding prevalence value. This produced a 1, 000 × 12, 000 binary matrix with an average of 3,534 genes per simulated genome.

In the well-specified case, we simulated an observed presence/absence data matrix (**X**) by multiplying the ground truth presence/absence data matrix (**Y**) by an observation matrix (**Z**). We drew a vector of 1,000 genome completeness values *λ* from a uniform distribution ranging from 0.05 to 1.1 and clipped values between 0 and 1, resulting in 99 complete simulated genomes and 901 incomplete simulated genomes ranging from 5% to 100% complete. We then generated the observation data matrix **Z** as a second set of Bernoulli random draws parameterized by the *λ* values. To generate the simulated observed presence/absence data, we took the entrywise (Hadamard) product of the ground truth presence/absence matrix **Y** and the observation matrix **Z**. Finally, any simulated genes without at least one detected presence were dropped from the dataset, resulting in a total of 9,594 gene families, of which 915 were core and 8,679 were flexible, including 1,485 extended core genes, 3,908 shell genes, and 4,201 cloud genes.

#### Inaccurate Genome Completeness Estimates

We simulated the effect of inaccurate genome completeness estimates on gene prevalence estimates using the same simulated dataset used in the well-specified case. However, instead of the true underlying genome completeness (*λ*) values, we supplied estimators with simulated noisy completeness estimates. Completeness is often calculated using subsets of single copy core marker genes, and there are currently 1,128 single copy core marker genes available for *Prochlorococcus* in the latest version of the benchmarking sets of universal single-copy orthologs database (BUSCO v6.1.0) [29]. We simulated the effect of using different numbers of single copy marker genes to calculate completeness, with *n*_Marker_ = 1, 5, 10, 50, 100, 500, 1000, 1128, 2000, 3000. For each marker gene set size, we simulated the number of marker genes observed for each genome by sampling from a binomial distribution with *n*_Marker_ trials and probability of success *λ_i_*. The noisy completeness estimates were then calculated as the number of marker genes observed divided by the marker gene set size.

#### Biases in Gene Detection

We evaluated the effect of variable detection rates between gene families by skewing the observation matrix (**Z**) using gene-specific detection biases. Gene detection biases were randomly drawn from Beta distributions with *α* = 1 and *β* = 1, 0.3, 0.1, 0.03, 0.01, 0.003, 0.001, 0.0003, 0.0001, with each draw producing a vector of gene detection biases, **b**, simulating different levels of variability in gene detection. For each simulated genome, the gene bias vector was used to produce a genome-specific vector of observation probabilities, **P***_i_*, adjusted so that, on average, the number of detected genes would approximate the product of the total number of genes in the corresponding simulated genome and the genome completeness parameter *λ_i_*. This was achieved by solving

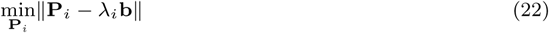

subject to 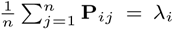 and 0 ≤ **P***_ij_* ≤ 1. The observation matrix **Z** was then generated as independent Bernoulli random draws, with the probability of observing gene *j* in genome *i* being equal to **P***_ij_*. The simulated observed presence/absence data matrix **X** was then generated in the same way as the unbiased simulation, taking the Hadamard product of matrices **Y** and **Z**, and removing any genes that were not detected at least once in the simulated presence/absence matrix.

#### Multicopy Genes

We evaluated the effect of multicopy genes on our gene prevalence estimation model by simulating gene presence/absence data generated from genomes containing multiple copies of genes. We tested a range of scenarios with overall mean gene copy numbers ranging from 1 (single copy) to 11. For each scenario, we designated a per-gene copy number parameter *e_j_*, where *e_j_* was randomly drawn from an exponential distribution parameterized with one of the following scaling factors: [0.001, 0.01, 0.023, 0.05, 0.1, 0.2, 0.5, 1, 2, 5, 10]. For each gene marked as present in the true presence/absence matrix **Y** (i.e. **Y***_ij_* = 1), we set the copy number to 1 + *c* where *c* was randomly drawn from a per-gene Poisson distribution parameterized by *e_j_*. This resulted in count matrix **C** where **C***_ij_* = 0 if **Y***_ij_* = 0 and **C***_ij_* ≥ 1 if **Y***_ij_* = 1. The observed presence/absence matrix **X** was generated by independently sampling the number of observed copies of each gene in each genome from a binomial distribution with **C***_ij_* trials and success probability *λ_i_*. This corresponds to independently observing each gene copy with probability equal to the simulated completeness of its genome, preserving genome completeness in expectation while allowing the realized fraction of observed copies to vary. Each entry **X***_ij_* was set to 1 if at least one copy of the corresponding gene was observed, and to 0 otherwise.

### Implementation

All model estimation procedures, confidence interval calculations, and simulated datasets were implemented in a publicly-available Python package. The library can be accessed via the public GitHub repository at https://github.com/blasks/gene-prevalence. The implementation follows conventions of the ‘scikit-learn’ library, with estimation and confidence interval methods accessible via the ‘BernoulliPrevalenceModel’ class found under the ‘model’ module of the package. The module ‘simulations’ contains code for constructing simulated datasets. All functions and classes contain complete document strings, and instructions for installing the library can be found in the project file ‘README.md’.

All experiments and calculations described in this manuscript were implemented in Jupyter notebooks and R scripts that can be found under the ‘experiments’ directory in the project repository. The code for generating the figures included in this manuscript can also be found in that directory.

#### *Prochlorococcus* Gene Presence/Absence Data

We generated gene presence/absence data from a dataset of 1,106 *Prochlorococcus* genomes collected as part of the ProSynTaxDB [31]. The dataset includes complete, closed genomes from culture isolates, as well as single-cell amplified genomes (SAGs) generated from single cell isolations collected from field sites throughout the global ocean. The clade membership of each genome was taken from Coe et al. (2025) based on their phylogenetic analysis. The genomes were downloaded from the Joint Genome Institute’s Integrated Microbial Genomes and Microbiomes database (IMG), along with their corresponding gene calls, functional annotations, and genome completeness estimates. The IMG annotation pipeline uses CheckM2 [25] to estimate genome completeness and based on the KEGG annotations of the genome [41]. We removed 7 genomes from the dataset for which genome completeness data was not available, leaving a total of 1,099 *Prochlorococcus* genomes (Supplementary File 1).

We used Anvi’o to generate presence/absence data from the *Prochlorococcus* genomes [42]. The genomes were imported into an Anvi’o genomes storage database using the ‘anvi-gen-contigs-database’ and ‘anvi-gen-genomes-storage’ commands. We then used the ‘anvi-pan-genome’ command to perform pangenome analysis on the dataset. Briefly, the analysis uses DIAMOND [43] to compute all-by-all amino acid sequence alignments followed by the the MCL algorithm [44] to cluster genes into families of orthologs. This produced a total of 52,500 unique orthologous gene family clusters, as well as the number of times each gene family occurs in each of the 1,099 genomes (Supplementary File 2). We further processed this data into a 1,099 x 52,500 matrix of binary (1/0) presence/absence data by applying a boolean mask to the count data, whereby 1 indicates a count ≥ 1 and 0 indicates a count of 0 for the corresponding gene in the corresponding genome.

#### Dataset Subsampling

Samples of the *Prochlorococcus* presence/absence data were constructed by using a pseudo random number generator to draw random rows from the matrix, representing a subset of genomes. For experiments comparing gene prevalence values between clades, samples were drawn exclusively from the rows corresponding to a particular clade (HLII and LLI in the experiments shown). For experiments comparing gene prevalence values between complete and incomplete genomes, samples were class-balanced by ensuring equal clade representation in the two samples. This was achieved by pre-determining the number of genomes to be drawn from each clade, and then constructing each sample as a combination of multiple clade subsamples.

#### Classifying *Prochlorococcus* ASVs by clade

We developed a pipeline to annotate the clade of the 616 amplicon sequence variants (ASVs) labeled as *Prochlorococcus* in the Global rRNA Universal Metabarcoding Plankton database (GRUMP) [32]. First, we assembled a database of reference amplicons using the 1,106 *Prochlorococcus* genomes included in the ProSynTaxDB [31]. Ribosomal 16S gene sequences were identified in each genome using Barrnap, imported into a QIIME 2 database [45], and trimmed using the universal primers 515Y and 926R that were used to amplify the sequences in the GRUMP database [32]. Incomplete sequences were removed from the database, resulting in 804 clade-labeled reference eASVs of 374 base pairs each. The clade of each environmental ASV was then inferred by searching ASVs against the reference database using the ‘classify-consensus-vsearch’ function of the QIIME 2 ‘feature-classifier’ module, using default settings and ‘top-hits-only’ set to true. This resulted in the successful classification of 586 of the ASVs. These were then filtered in a final step to exclude clade-level classifications that were not unique (i.e. cases where an ASV had top hits to multiple clades). This yielded a final set of 451 GRUMP ASVs with a predicted *Prochlorococcus* clade classification (Supplementary File 3). The remaining 165 ASVs with ambiguous or missing vsearch results were labeled as “unclassified”.

#### Estimating Global *Prochlorococcus* Gene Prevalence

To estimate gene prevalence in the global *Prochlorococcus* population, we first estimated the relative abundance of each *Prochlorococcus* clade using the clade-annotated ASVs from the GRUMP dataset [32]. GRUMP Samples were grouped according to Longhurst biome [33] and sampling depth, using three biomes (Coastal, Trades, and Westerlies) and three depth ranges (0–50 meters, 50–100 meters, and 100– 200 meters). The Polar biome and depths below 200 meters were excluded because *Prochlorococcus*is predominantly distributed in the euphotic zone of tropical and subtropical oceans, and generally absent from polar waters [5, 46]. Within each biome–depth bin, ASV relative abundances were aggregated to estimate the mean clade composition of the *Prochlorococcus* population (Supplementary File 4). ASVs that could not be unambiguously assigned to a clade were retained as an “unclassified” category. To evaluate whether this a priori stratification adequately captured reproducible variation in clade composition, we compared it against an unconstrained clustering approach based on agglomerative hierarchical clustering of sample-level clade profiles. Although the data-driven clusters produced greater within-group similarity, they explained only modestly more variance than the biome–depth bins, indicating that the ecologically defined bins captured much of the large-scale structure in the dataset. We therefore used the biome–depth bins for downstream analyses because they are ecologically interpretable and defined independently of the composition of the GRUMP dataset.

We next estimated the relative contribution of each biome–depth bin to the global *Prochlorococcus* population using an annual climatological model of *Prochlorococcus* cell abundance [46]. Cell concentrations were integrated over each depth interval and combined with Longhurst province surface areas to estimate the total number of *Prochlorococcus*cells represented by each biome–depth bin. The clade composition estimates from GRUMP were then multiplied by the corresponding bin-level cell abundance, summed across bins, and normalized to obtain a single estimate of the relative abundance of each clade in the global *Prochlorococcus* population.

We independently estimated gene prevalence within each clade using the maximum likelihood approach described above (Supplementary File 5). We then combined the resulting clade-specific estimates according to the estimated global relative abundance of each clade. Prevalence estimates corresponding to the unclassified ASV fraction were generated using all 1,099 reference genomes. Global prevalence and confidence intervals were calculated using the weighted-combination procedures described above. Gene families were categorized using the prevalence thresholds defined by [13]. In particular, gene families with prevalence ≤ 0.15 were classified as cloud genes, those with prevalence *>* 0.15 and *<* 0.9 as shell genes, and those with prevalence ≥ 0.9 as extended core genes. To define core genes while allowing for uncertainty introduced by sequencing, assembly, and gene calling, we classified a gene family as core when its 95% confidence interval included a prevalence of 1. The combined global prevalence estimate for each of the 52,500 gene families in the analyzed *Prochlorococcus* pangenome can be found in Supplementary File 6, along with confidence intervals, gene category, and the most frequent KEGG annotation for each gene family.

## Conflicts of Interest

The authors declare that they have no competing interests.

## Funding

This work was supported by a grant from the Simons Foundation to E. V. A. (Award ID 549945FY22) and to J.B. (Award ID 54993FY22). Y.R. was supported through funding from Simons Collaboration on Computational Biogeochemical Modeling of Marine Ecosystems (Grant ID: 549931 to Michael J. Follows).

## Data and Code Availability

All genomes, annotations, and genome completeness data is accessible via the Joint Genome Institute’s Integrated Microbial Genomes and Metagenomes database (IMG/M), using the accession numbers listed in Supplementary File 1. Global 16S rRNA data from the GRUMP database [32] is accessible on Zenodo (https://doi.org/10.5281/zenodo.15586714). Estimated climatology of global *Prochlorococcus* cell counts [46] are available through the Marine Biological Laboratory (MBL) and Woods Hole Oceanographic Institution (WHOI) library (https://hdl.handle.net/1912/25779). Longhurst province boundaries were downloaded from marineregions.org (https://www.marineregions.org/sources.php#longhurst). All code used to run experiments and generate figures is available as part of the project repository (https://github.com/blasks/gene-prevalence).

## Generative AI Disclosure

Codex (GPT-5.4 through GPT-6), and Claude Code (Sonnet 4.6, Sonnet 5, Opus 4.8, and Opus 5) were used to help generate, modify, and debug code used in this manuscript; all AI-generated code was reviewed and tested by one or more authors. Chat GPT-5.6 was used to review the manuscript for typos and grammatical errors. Authors take responsibility for the integrity of all content in this manuscript.

## Author Contributions Statement

Conceptualization: S.B. and E.V.A. Data curation: S.B. Formal analysis: S.B. and L.G.C. Funding acquisition: E.V.A., J.B., and J.A.F. Investigation: S.B. and L.G.C. Methodology: S.B., L.G.C., and J.B. Project administration: E.V.A. Software: S.B. and L.G.C. Supervision: E.V.A. and J.B. Validation: S.B., L.G.C., and Y.R. Visualization: S.B., L.G.C., and Y.R. Writing—original draft: S.B. and L.G.C. Writing—review and editing: S.B., L.G.C., Y.R., J.A.F., J.B., and E.V.A.

## Acknowledgments

The authors thank their colleagues in the Simons Collaboration on Computational Biogeochemical Modeling of Marine Ecosystems (CBIOMES), and especially the members of the Armbrust lab for helpful discussions and insights.

## Supplementary Figures

**Supplementary Figure 1.**
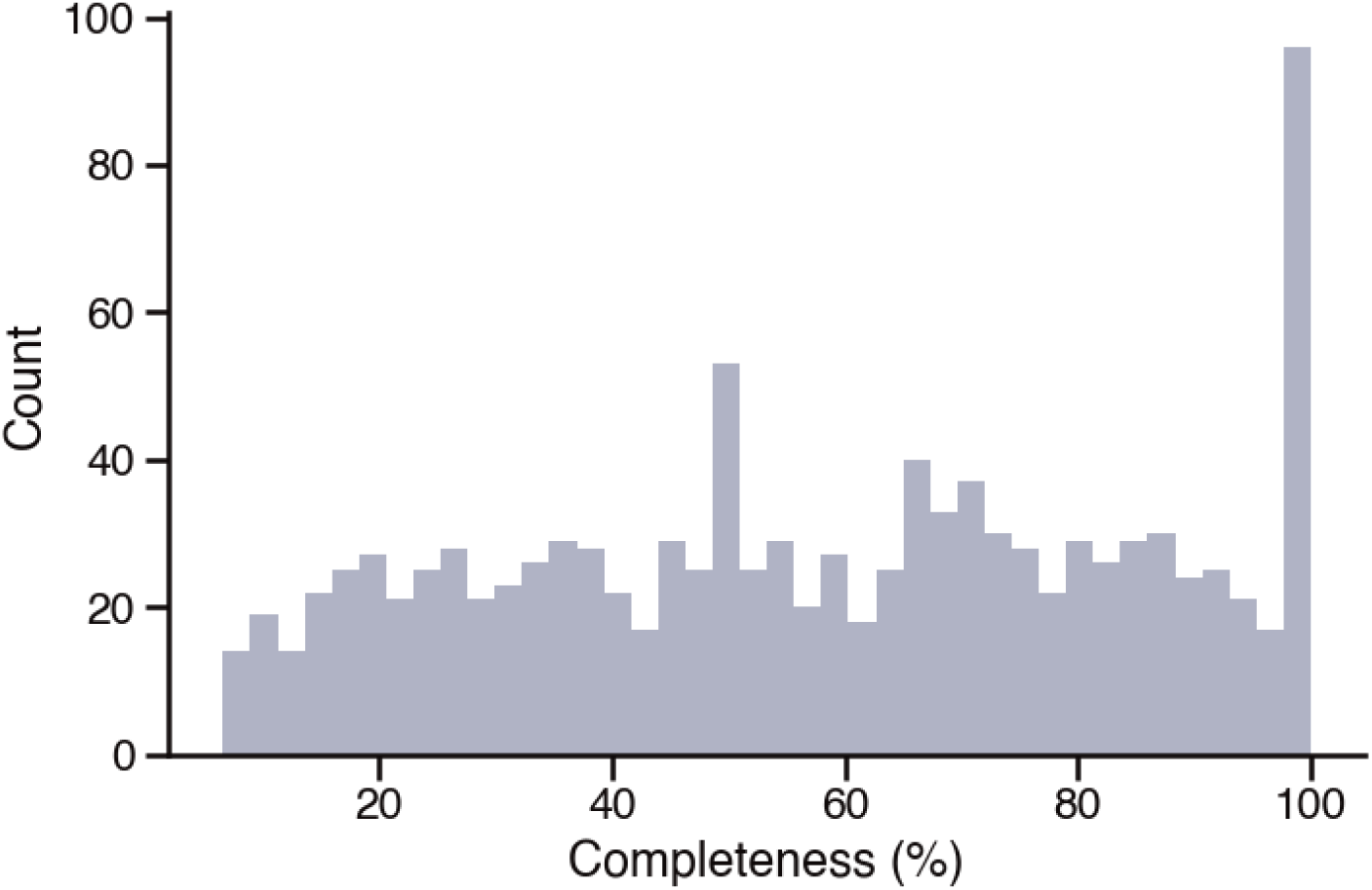
Out of 1,099 available *Prochlorococcus* genomes, 1,034 are incomplete. Distribution of genome completeness values of *Prochlorococcus* genomes, as estimated by CheckM2.

**Supplementary Figure 2.**
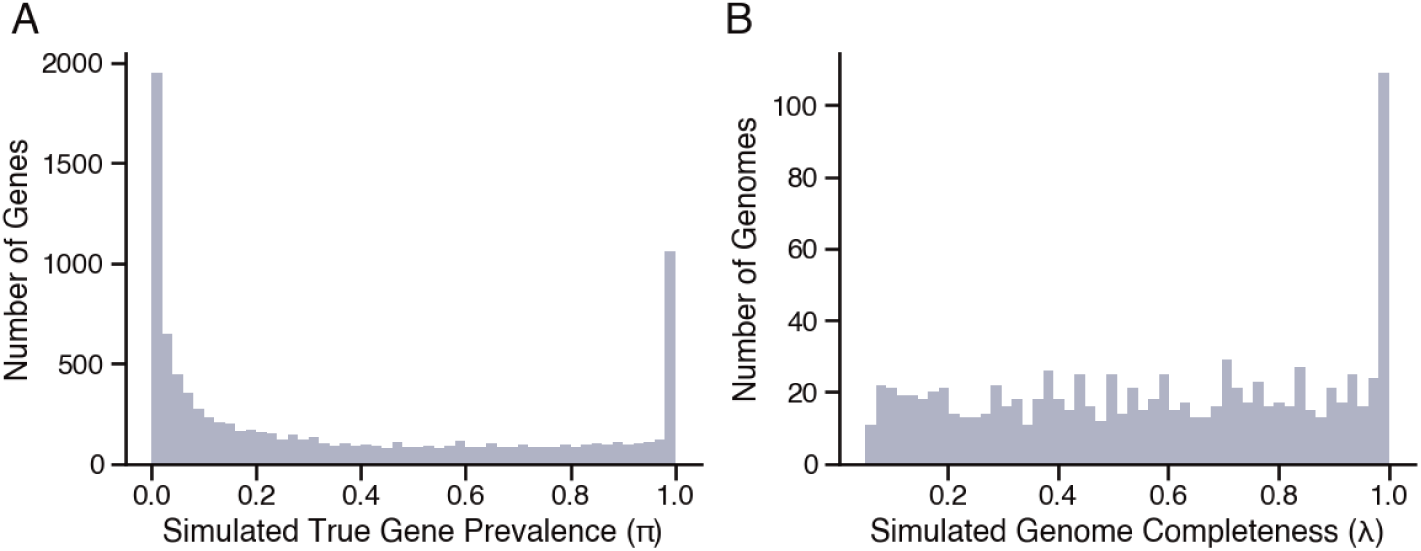
Simulated presence/absence data was generated from ground truth parameters drawn from parametric distributions. A) Distribution of true prevalence values for 9,594 simulated gene families, drawn from Beta distribution, of which 915 were core (prevalence = 1.0). B) Distribution of completeness values for 1,000 simulated genomes, drawn from uniform distribution with 99 genomes set to complete.

**Supplementary Figure 3.**
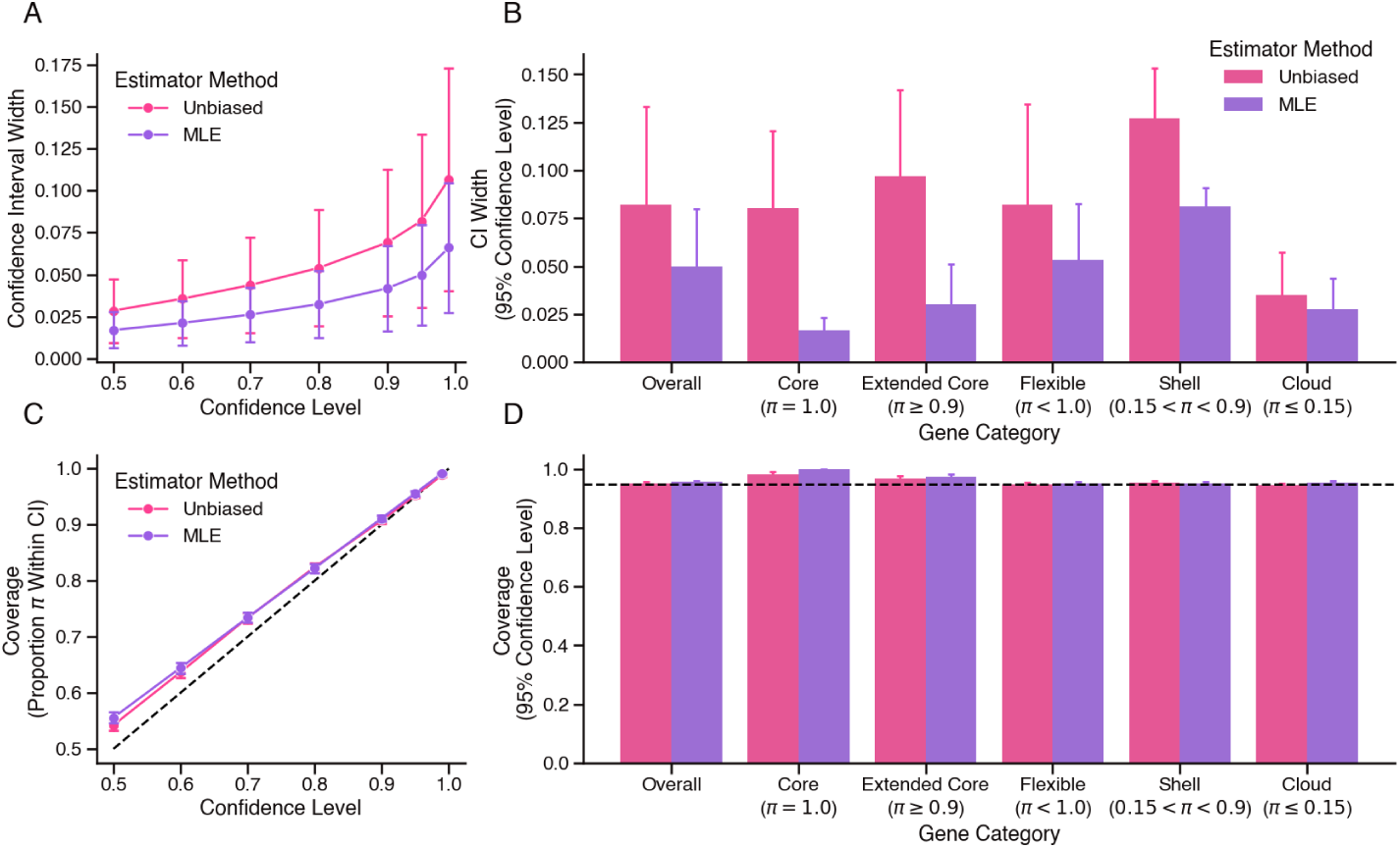
Simulation experiments show narrower, well-calibrated confidence intervals using likelihood ratio methodology in combination with MLE estimator, as compared to Gaussian approximation methodology in combination with unbiased estimator. A) Mean width of Confidence Intervals (CIs) of method estimates as a function of confidence level (1 − *α*). Bars indicate standard deviation. B) Mean CI width at 95% confidence level, broken down by gene subset. Bars indicate standard deviation. C) Coverage (proportion of genes for which prevalence falls within CI) as a function of confidence level (1 − *α*). D) Coverage at 95% confidence level, broken down by gene subset. Bars indicate 2× standard error.

**Supplementary Figure 4.**
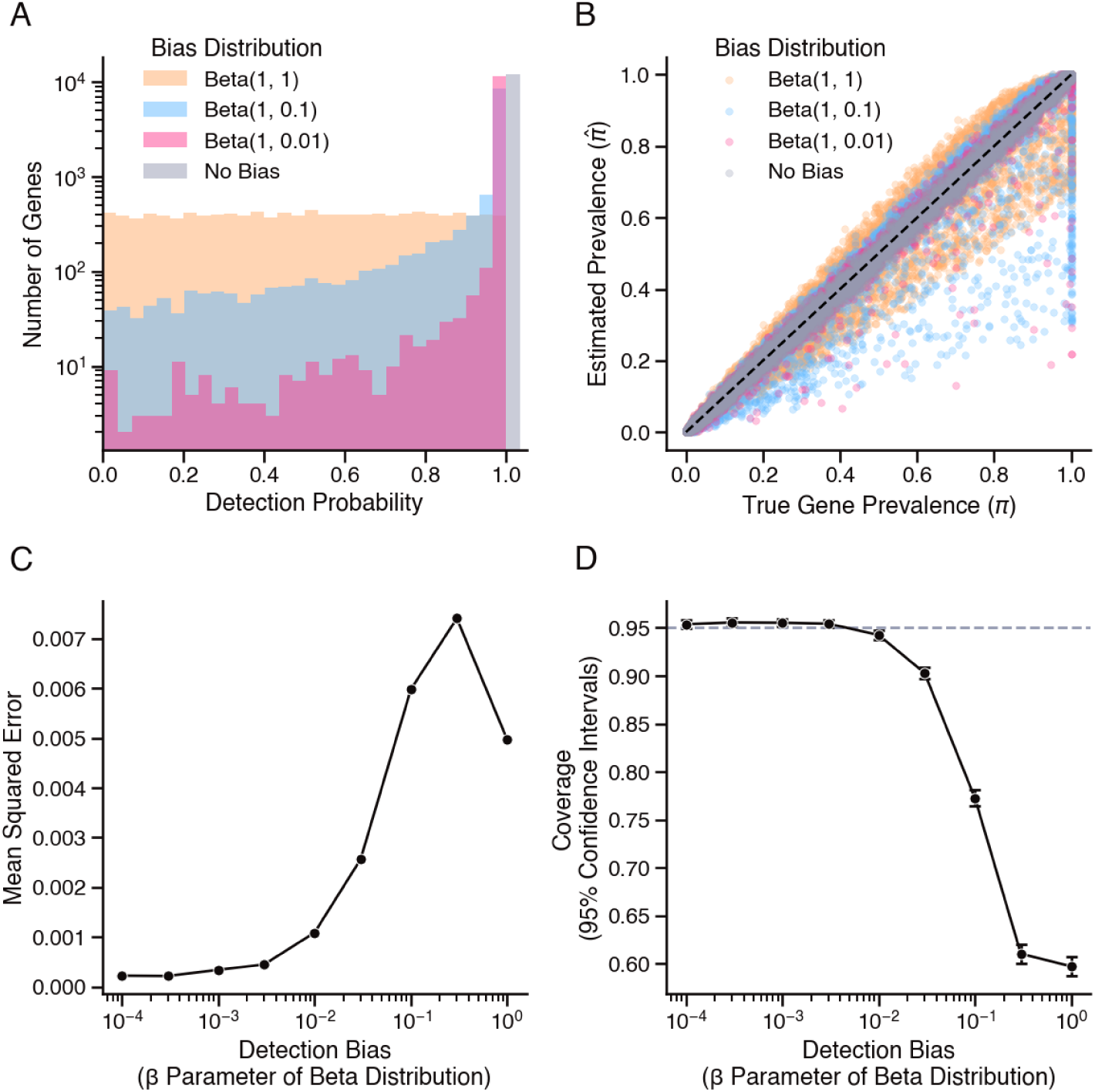
Simulation experiments show more variation in gene detection rates results in higher error and empirical coverage below target confidence level. A) Distributions of simulated gene detection probabilities generated from Beta distributions. B) Estimated versus true gene prevalence from models fit to simulated presence/absence data, generated by taking into account gene detection biases. Black dotted line indicates perfect agreement. C) Mean squared error and D) coverage from model estimates fit to simulated data with increasing bias in gene detection rates. Bars indicate 2× standard error.

**Supplementary Figure 5.**
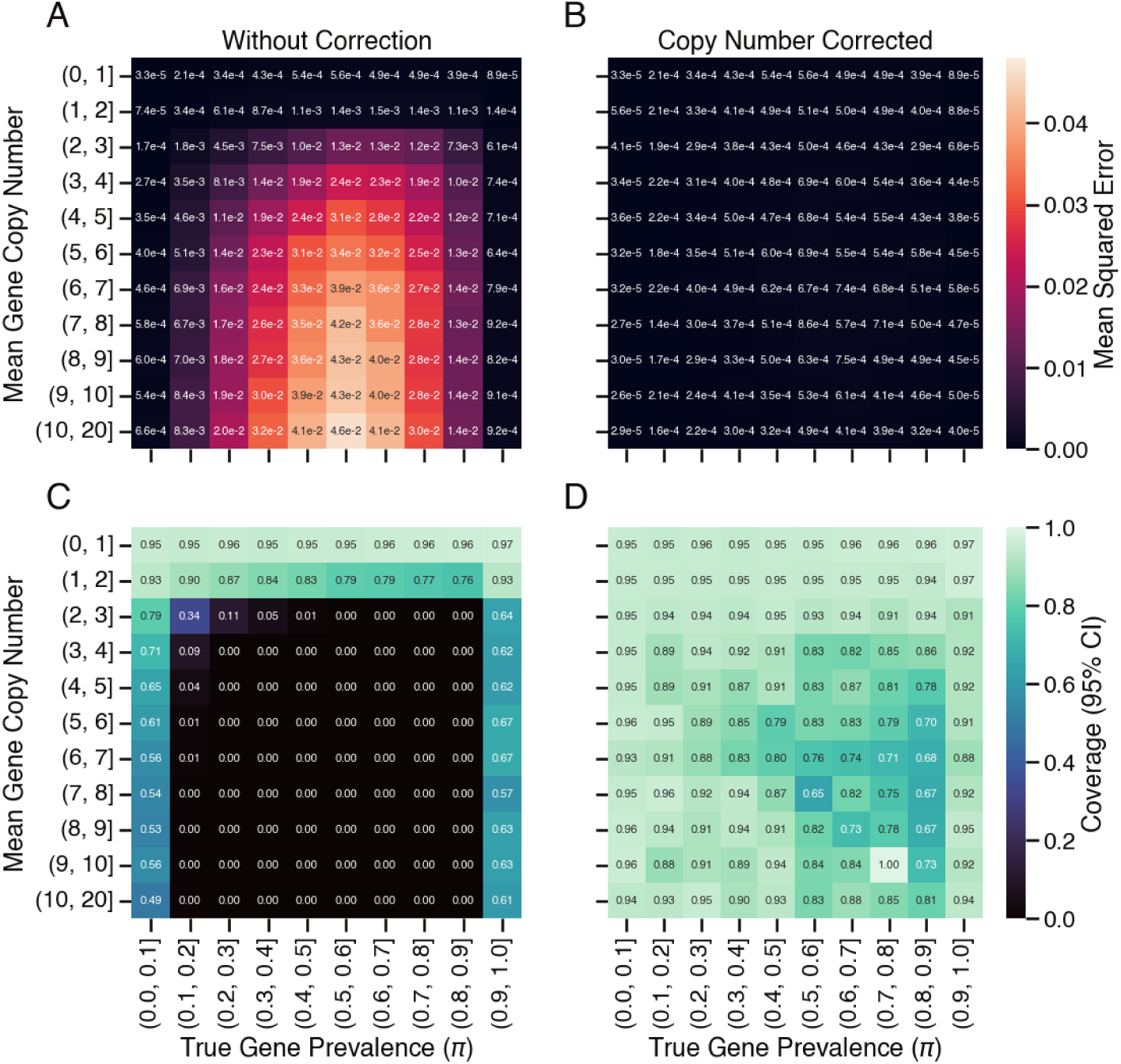
In the absence of copy number correction, method performance on simulated datasets is worse in gene families with intermediate prevalence and a mean copy number above 2. A) Mean squared error (MSE) of maximum likelihood estimation (MLE) without copy number correction, binned by true underlying gene family prevalence (x-axis) and true mean gene family copy number (y-axis), calculated prior to masking incomplete genomes. B) MSE of copy-number-corrected MLE, binned by gene family prevalence and mean copy number. C) Coverage (proportion of genes for which prevalence falls within confidence interval) at 95% confidence level, for MLE without copy number correction, binned by gene family prevalence and mean copy number. D) Coverage for copy-number-corrected MLE, binned by gene family prevalence and mean copy number.

**Supplementary Figure 6.**
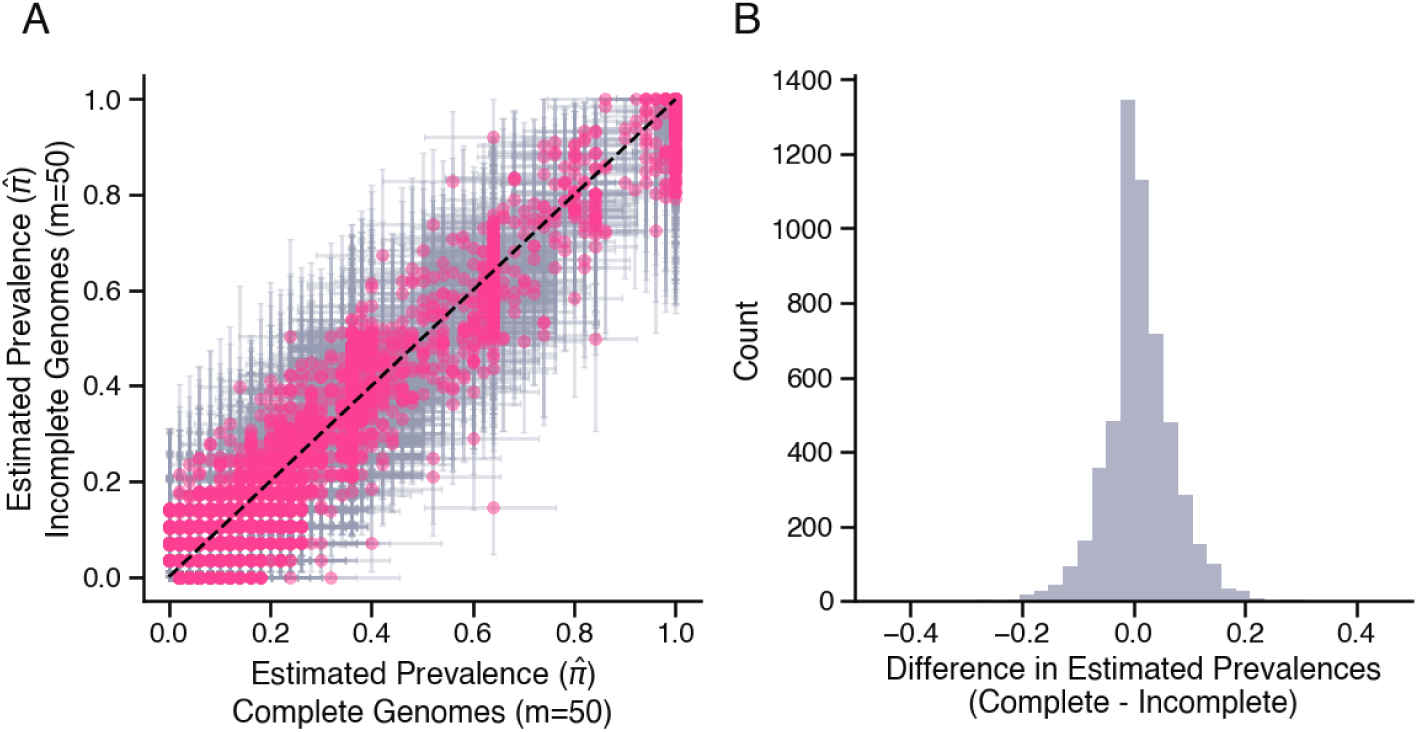
Comparison of model outputs from subset of complete and incomplete genomes suggests real data bias is well-controlled by model assumptions. A) Comparison of gene prevalence estimates from models fit to subsamples (*m* = 50, clade-balanced) of complete (x-axis) and incomplete (y-axis) *Prochlorococcus* genomes. Bars indicate 95% confidence intervals and black dotted line indicates perfect agreement. B) Histogram of differences in prevalence estimates from the two models fit to genome subsamples. Zero values have been removed for ease of visualization.

**Supplementary Figure 7.**
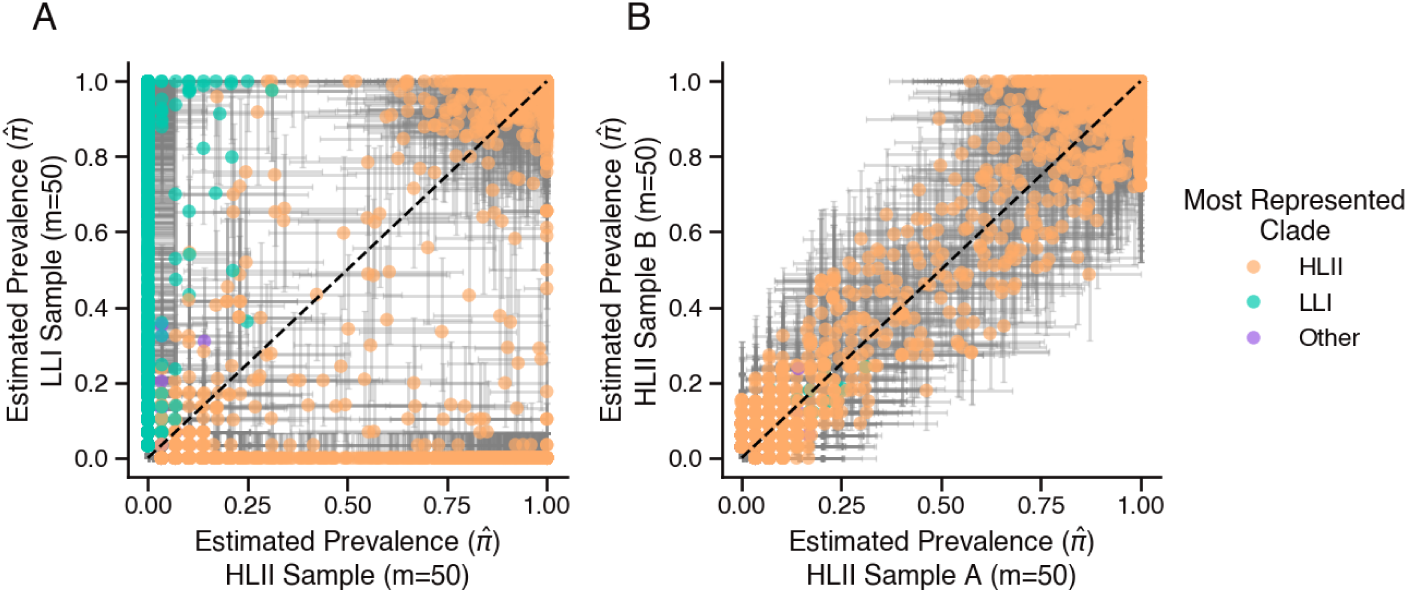
Prevalence estimates exhibit different distributions between clade subpopulations, but are consistent between independent subsamples from within the same clade. A) Comparison of gene prevalence estimates from models fit to subsamples (*m* = 50) of *Prochlorococcus* genomes from the High Light II (HLII) clade (x-axis) and the Low Light I (LLI) clade (y-axis). B) Comparison of gene prevalence estimates from models fit to independent, non-overlapping subsamples (*m* = 50) of genomes from the HLII clade. Bars indicate 95% confidence intervals and black dotted lines indicate perfect agreement.

**Supplementary Figure 8.**
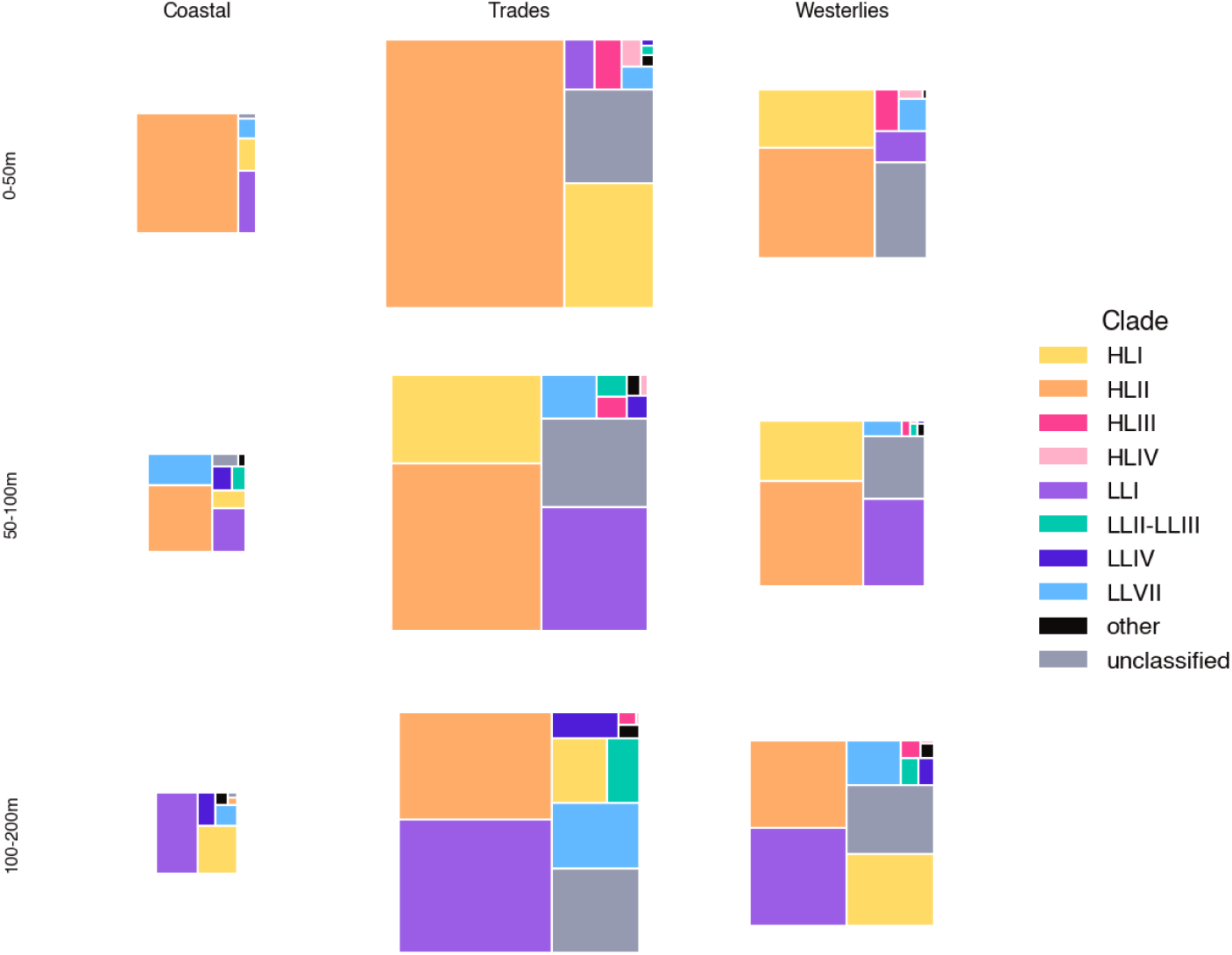
Relative abundance of *Prochlorococcus* clades varies by ocean biome and depth. Color indicates clade membership and size indicates relative difference in mean annual population.

**Supplementary Figure 9.**
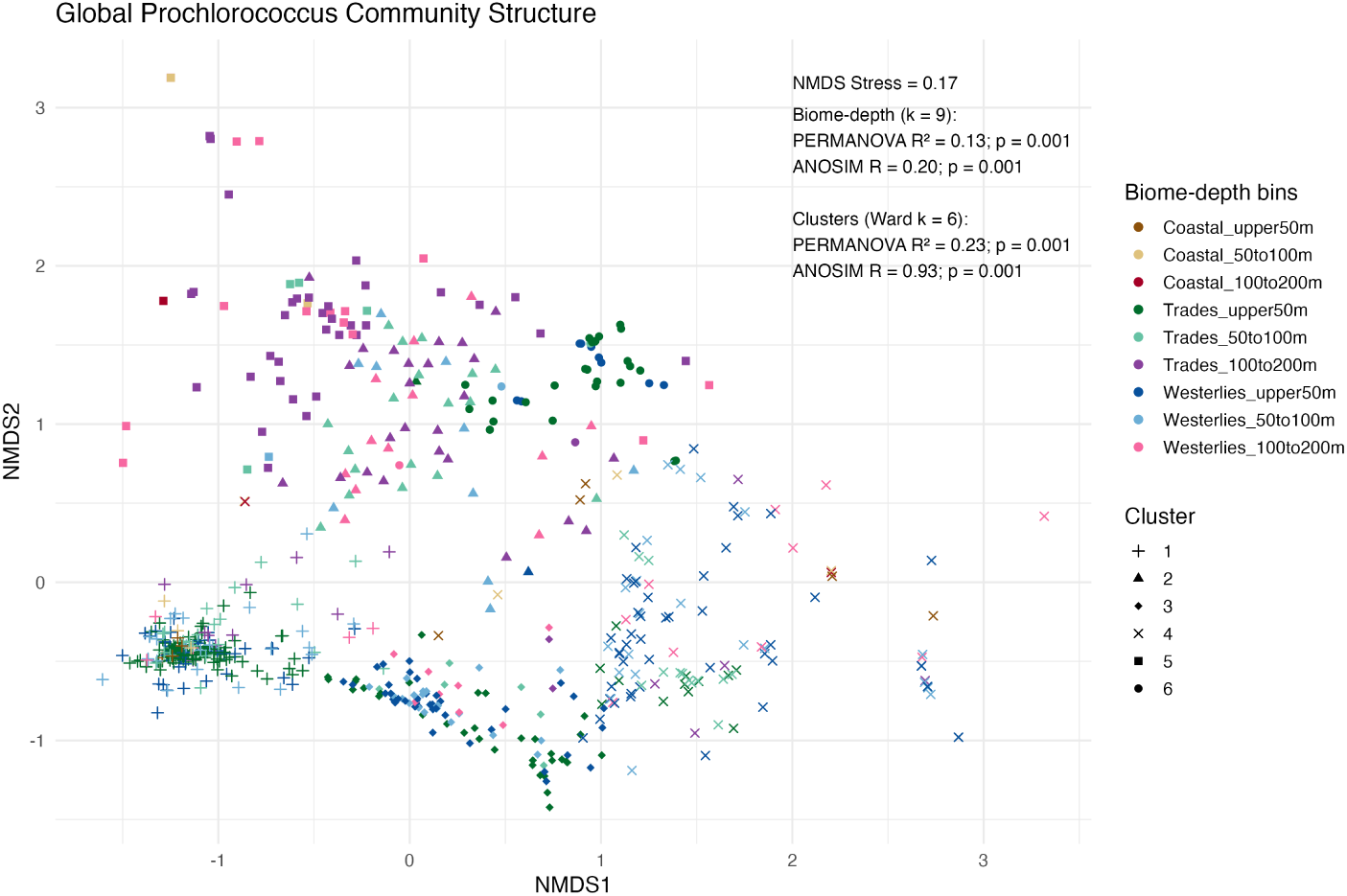
Comparison of unsupervised versus a-priori clustering of samples from global rRNA amplicon sequence variant database (GRUMP) [32] indicates *Prochlorococcus* community structure is largely governed by broad environmental gradients rather than discrete community states. NMDS plot of clade compositions of *Prochlorococcus* reads from GRUMP samples with color indicating a priori biome–depths bin membership (k=9) and shape indicating hierarchical cluster membership (k=6).

**Supplementary Figure 10.**
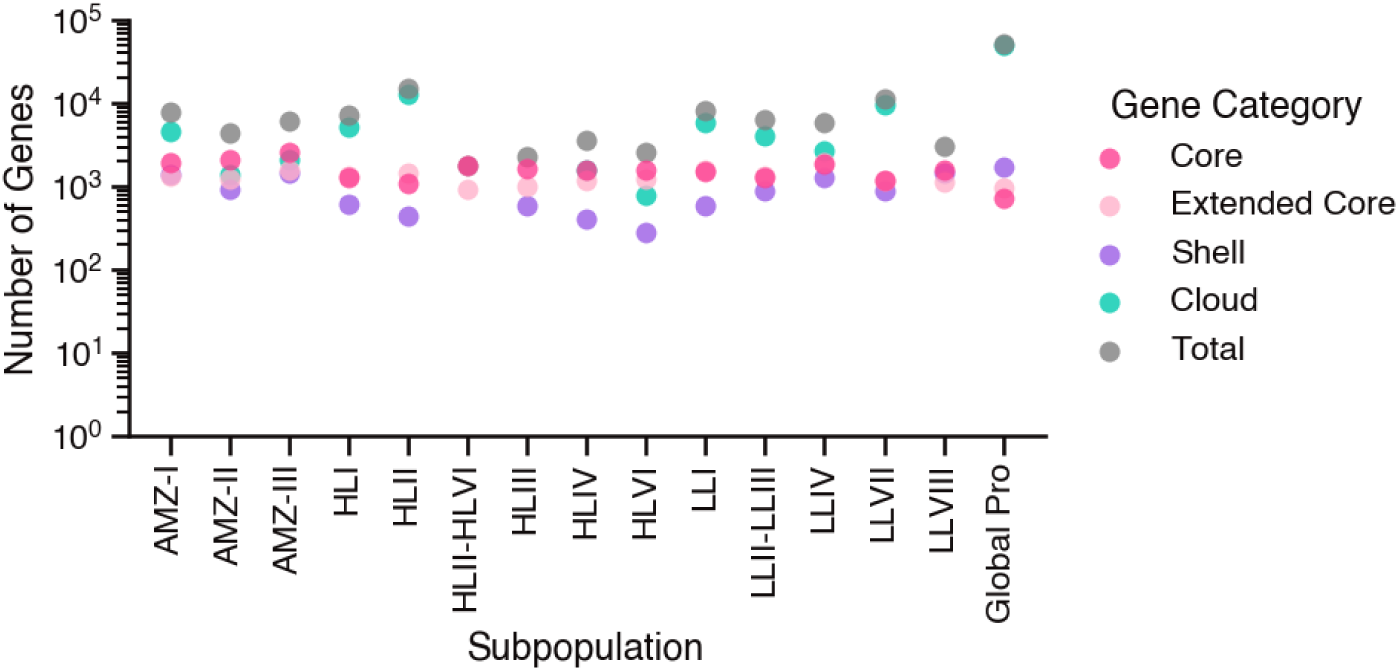
Number of genes core to global *Prochlorococcus* population is less than the number of genes core to any given *Prochlorococcus* clade. Color indicates pangenome category of genes and bar height indicates number of genes in the category.

## Descriptions of Additional Supplementary Files

**Supplementary File 1**

Metadata for 1,099 *Prochlorococcus* genomes used in this study. Sequences and annotations can be retrieved from the Joint Genome Institute’s Integrated Microbial Genomes and Metagenomes database (IMG/M) using the accession number under field “IMG Genome ID”. CheckM2 completeness score can be found under field “Completeness”.

**Supplementary File 2**

Gene count matrix for 1,099 genomes and 52,500 gene families of the analyzed *Prochlorococcus* pangenome. Each entry indicates the number of times the corresponding gene was detected in each genome.

**Supplementary File 3**

Clade classification for every 16S rRNA amplicon sequence variant (ASV) in the GRUMP data base [32] originating from *Prochlorococcus*.

**Supplementary File 4**

Relative abundance of *Prochlorococcus* clades in the global ocean, based on aggregated GRUMP ASV counts [32] and binned by Longhurst biome [33] and depth.

**Supplementary File 5**

Clade-specific gene prevalence estimates for every gene family in the detected *Prochlorococcus* pangenome.

**Supplementary File 6**

Overall global gene prevalence estimates for every gene family in the detected *Prochlorococcus* pangenome.

## Notes

### Competing Interest Statement

The authors have declared no competing interest.

